# Shelterin is a Dimeric Complex with Extensive Structural Heterogeneity

**DOI:** 10.1101/2022.01.24.477420

**Authors:** John C Zinder, Paul Dominic B Olinares, Vladimir Svetlov, Martin W Bush, Evgeny Nudler, Brian T Chait, Thomas Walz, Titia de Lange

## Abstract

Human shelterin is a six-subunit complex – comprised of TRF1, TRF2, Rap1, TIN2, TPP1, and POT1 – that binds telomeres, protects them from the DNA-damage response, and regulates the maintenance of telomeric DNA. Although high-resolution structures have been generated of the individual structured domains within shelterin, the architecture and stoichiometry of the full complex are currently unknown. Here we report the purification of shelterin subcomplexes and reconstitution of the entire complex using full-length, recombinantly produced components. By combining negative-stain electron microscopy (EM), crosslinking mass spectrometry (XLMS), mass photometry, and native mass spectrometry (MS), we obtain stoichiometries as well as domain-scale architectures of shelterin subcomplexes and determine that they are extensively conformationally heterogenous. For POT1/TPP1 and POT1/TPP1/TIN2, we observe high variability in the positioning of the POT1 DNA-binding domain, the TPP1 OB fold, and the TIN2 TRFH domain with respect to the C-terminal domains of POT1. Truncation of unstructured linker regions in TIN2, TPP1, and POT1 did not reduce the conformational variability of the heterotrimer. Both shelterin and the TRF1/TIN2/TPP1/POT1 subcomplex primarily adopt fully dimeric complexes, even in the absence of DNA substrates. TRF1/TIN2/TPP1/POT1 and shelterin complex showed extensive conformational variability, regardless of the presence of DNA substrates. We conclude that shelterin adopts a multitude of conformations and argue that its unusual architectural variability is beneficial for its many functions at telomeres.

## Introduction

Telomeres are protective assemblies of DNA and proteins at the ends of linear chromosomes. Human telomeres are composed of the hexanucleotide 5’TTAGGG3’ sequence repeated for 3-10 kilobases and terminate in a ~100-nucleotide 3’ single-stranded (ss) overhang (de Lange, 2018). Telomeres need to solve two problems 1) The DNA-replication machinery is unable to replicate all the way to the end, resulting in terminal attrition as cells divide (‘The end-replication problem’) and 2) Telomere ends resemble sites of DNA damage that must be protected against erroneous repair and inappropriate activation of DNA-damage checkpoints (‘The end-protection problem’). The telomerase ribonucleoprotein extends telomeres, using its RNA component as a template for its reverse-transcriptase activity, to solve the end-replication problem (Greider & Blackburn, 1985; Lingner et al., 1997). Shelterin, a multi-subunit DNA-binding complex (de Lange, 2005), solves the end-protection problem and additionally recruits telomerase to solve the end-replication problem (Nandakumar et al., 2012; Zhong et al., 2012). Defects in shelterin components are associated with diverse pathologies including familial cancer predisposition and premature aging, underscoring its importance in cellular physiology and development (Maciejowski & de Lange, 2017; Artandi & DePinho, 2010; Haycock et al., 2017; Nelson & Bertuch, 2012; Savage et al., 2011; Savage & Bertuch, 2010).

Shelterin specifically binds to double-stranded (ds) telomeric repeat sequences via the C-terminal Myb domains of its TRF1 and TRF2 subunits (Chong et al., 1995; Broccoli et al., 1997; König et al., 1998; Bilaud et al., 1997). TRF1 and TRF2 homodimerize via their TRF Homology (TRFH) domains (Bianchi et al., 1997; Fairall et al., 2001) and are bridged by TIN2 (Ye et al., 2004a; Liu et al., 2004a; Kim et al., 1999), which contains a variant of a TRFH domain that does not dimerize (Hu et al., 2017). In somatic cells, TRF2 is critical for the formation of t-loops (Griffith et al., 1999; Doksani et al., 2013; Timashev & de Lange, 2020), protective structures in which the telomeric 3’ overhang invades into the dsDNA region, hiding the 3’ end from recognition by the ATM kinase DNA-damage response (DDR) pathway and double-strand break repair by classical non-homologous end joining (de Lange, 2018). TIN2 is recruited to telomeres primarily via its interaction with TRF1, which is stronger than its interaction with TRF2 (Ye et al., 2004a; Frescas & de Lange, 2014; Takai et al., 2011; Hu et al., 2017; Chen et al., 2008). TIN2 additionally interacts with TPP1 (Ye et al., 2004b; Liu et al., 2004b; Houghtaling et al., 2004), which in turn binds POT1, an ssDNA-binding protein that prevents replication protein A from accumulating at telomeric ssDNA and activating the ATR kinase DDR pathway (Baumann & Cech, 2001; Hockemeyer et al., 2006; Denchi & de Lange, 2007; Gong & de Lange, 2010; Kratz & de Lange, 2018; Wu et al., 2006). Finally, Rap1 binds tightly to TRF2, but is not required TRF2’s primary functions at telomeres (Timashev & de Lange, 2020; Sfeir et al., 2010; Martínez et al., 2013).

Shelterin acts in conjunction with several accessory factors such as the BLM and RTEL helicases, which facilitate the replication of telomeric DNA (Sfeir et al., 2009; Sarek et al., 2015; Zimmermann et al., 2014; Yang et al., 2020). After replication, shelterin ensures the reconstitution of the correct structure of the telomere end by recruiting the Apollo nuclease (via TRF2) (van Overbeek & de Lange, 2006; Lenain et al., 2006; Wu et al., 2010; Lam et al., 2010) and the CST fill-in machinery (via TPP1/POT1) to telomeres (Wu et al., 2012; Wan et al., 2009). Finally, shelterin regulates telomerase-mediated telomere extension by recruiting the enzyme via TPP1 (Nandakumar et al., 2012; Zhong et al., 2012) and limiting its activity at the longest telomeres (via TPP1/POT1-dependent recruitment of CST) (Chen et al., 2012).

All six shelterin components contain structured domains separated by linker regions that are predicted to be largely disordered (de Lange, 2018; Hu et al., 2017; Lim & Cech, 2021). These regions range in length from 25 to 150 amino acids (aa) in humans and, while their sequences and lengths are highly variable in other organisms, the trend of alternating structured domains and regions of predicted disorder is maintained in shelterin components throughout metazoan evolution (Myler et al., 2021). X-ray crystallography and NMR spectroscopy have generated molecular models of all known structured domains and minimal interacting motifs within human shelterin (Fairall et al., 2001; Court et al., 2005; Hu et al., 2017; Rice et al., 2017; Chen et al., 2017; Wang et al., 2007; Lei et al., 2004; Chen et al., 2011; Chen et al., 2008; Hanaoka et al., 2001), but there are currently no high-resolution structural models for full-length shelterin proteins. It has been speculated that shelterin is too flexible to be structurally characterized at high resolution (Lim & Cech, 2021), although experimental evidence for this notion is lacking.

Purification of mouse shelterin overexpressed in HEK293 cells has been reported, but that study did not yield sufficient quantities to probe its structure or stoichiometry (Erdel et al., 2017). In another study, a human shelterin subcomplex lacking TRF1 was purified from insect cells and a POT1:TPP1:TIN2:TRF2:Rap1 stoichiometry of 1:1:1:2:2 was proposed based on in-gel densitometry measurements of labeled components and size-exclusion chromatography coupled to multiple-angle light scattering, but no structural characterization of this complex was reported (Lim et al., 2017).

Here, we report single-particle negative-stain EM characterization of purified, recombinantly produced human shelterin subcomplexes containing full-length components and the reconstituted full, six-subunit complex. We employ streptavidin-DNA labeling and XLMS to identify elements within reference-free 2D-class averages of shelterin subcomplexes and arrive at domain-scale models of their structures in different conformations. Finally, we use mass photometry and native mass spectrometry to reveal that either one or two TIN2/TPP1/POT1 trimers can bind the TRF1 homodimer in reconstituted complexes, arguing that shelterin can exist as a fully dimeric complex. Mass photometry of reconstituted shelterin revealed that it could adopt a fully dimeric stoichiometry as well. The data reveal an extraordinary level of structural variability for shelterin and several of its subcomplexes. We discuss the implications of this variability in the context of the many different functions of shelterin.

## Results

### High Variability in the Relative Positioning of the DNA-Binding and C-terminal Domains of POT1

We first sought to structurally characterize a small, well-defined subcomplex of shelterin containing full-length POT1 and the N-terminal domain (comprising the POT1-binding domain and the OB fold) of TPP1 (POT1/TPP1^N^; Figure 1A). Recombinantly expressed POT1/TPP1^N^ could be purified to homogeneity as assessed by SDS-PAGE (Figure 1B) and bound to telomeric ssDNA with high specificity (Figure 1C).

**Figure 1.**
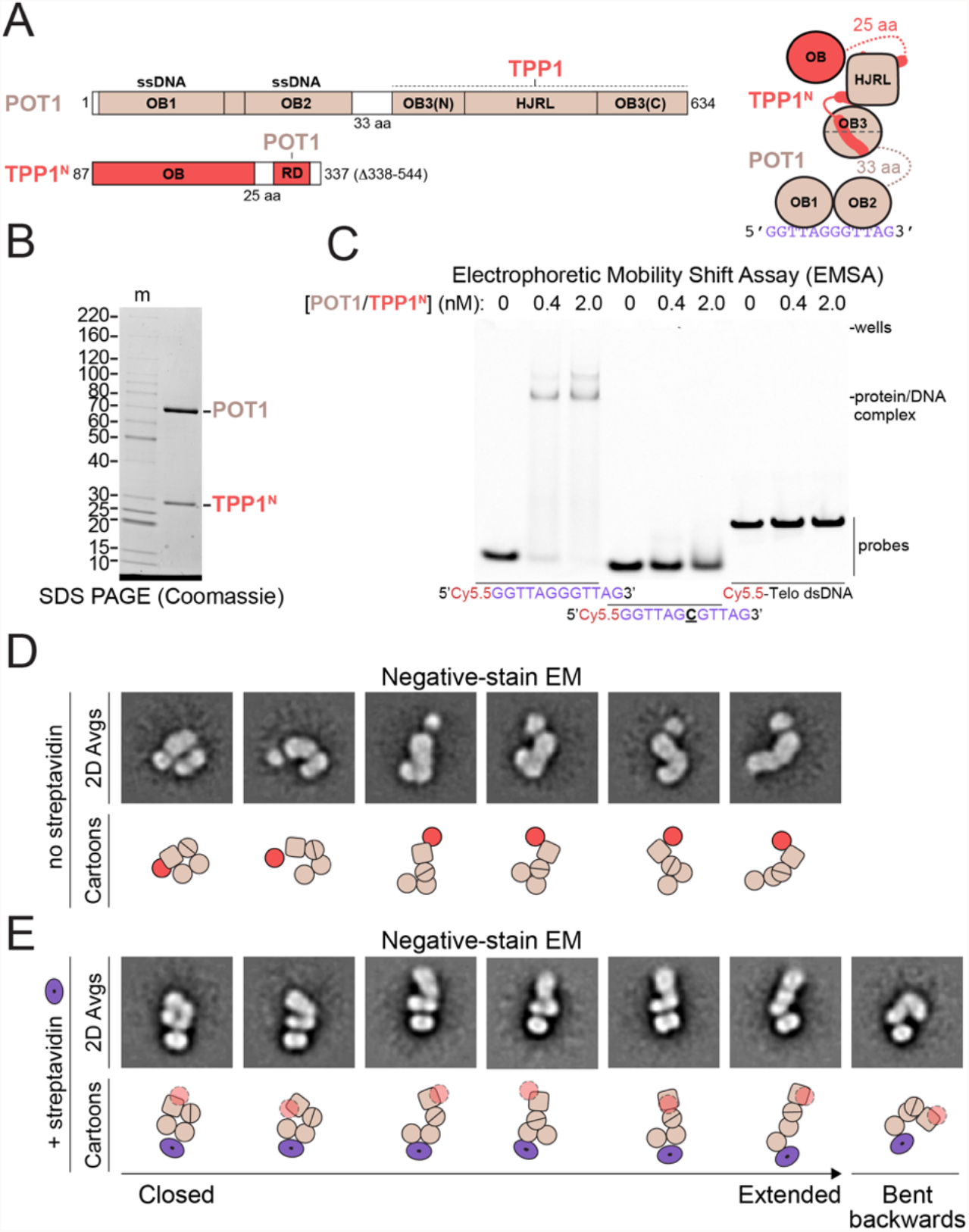
Structural variability and DNA binding of POT1/TPP1^N^. **(A)** Domain and cartoon schematics for POT1 and TPP1^N^ with starting and ending amino acids and flexible linker lengths indicated. Regions that associate with other proteins in shelterin are indicated. RD: recruitment domain; OB: oligonucleotide/oligosaccharide-binding domain; HJRL: Holliday junction resolvase-like domain. Regions of known structure are depicted in color in the domain schematics and as shapes in the cartoon. Regions of unknown structure are depicted in white in the domain schematic and as dashed lines with amino acid lengths indicated. **(B)** SDS-PAGE of purified POT1/TPP1^N^. Gel is 8-16% polyacrylamide Tris-glycine and stained with Coomassie blue. Molecular-weight markers (m) are shown in kDa. **(C)** DNA-binding activity of POT1/TPP1^N^ on fluorescent DNAs. Protein concentrations and DNAs are indicated. Probe concentration is 0.25 nM. Gel is 4-20% polyacrylamide TBE and was imaged for Cy5.5 fluorescence. **(D)** Negative-stain EM analysis of POT1/TPP1^N^ bound to 5’Biotin(GGTTAG)_2_ DNA. Reference-free 2D-class averages showing three lobes in a range of conformations were selected for display with cartoon interpretations below. Domains are as in panel A. **(E)** Negative-stain EM analysis of POT1/TPP1^N^ bound to streptavidin-5’Biotin(GGTTAG)_2_ DNA. Reference-free 2D-class averages showing a bipartite lobe associated with streptavidin and a second oblong lobe in a range of conformations were selected for display with cartoon interpretations below. Domains in the cartoons are as in panel A with TPP1’s OB fold depicted as transparent because its position is not discernable. The streptavidin tetramer is shown as a purple torus.

Negative-stain EM images of POT1/TPP1^N^ bound to ssDNA showed particles of similar size but with highly variable shapes (Figure 1—figure supplement 1A). Approximately 60,000 particles were auto-picked from 100 micrographs and, after centering and curation to 40,000 particles, subjected to 2D classification using the “Iterative Stable Alignment and Clustering” (ISAC) algorithm, an approach optimized for analyzing heterogenous samples (Yang et al., 2012). Details for this and other negative-stain EM datasets are provided in Supplemental Table 1.

Many of the reference-free 2D-class averages of POT1/TPP1^N^ showed an oblong lobe connected on one side to another oblong, often bipartite, lobe of similar size, and on the other side to a third, smaller and more circular lobe (Figure 1D; Figure 1—figure supplement 1A). The conformations of the two oblong lobes relative to each other ranged from almost fully extended to entirely closed. This finding suggests that the 33-aa linker connecting the N- and C-terminal domains of POT1 (Figure 1A) allows the two domains to freely move with respect to each other. The smaller, more circular lobe is likely the OB fold of TPP1, which is present in many positions relative to the POT1 C-terminal domain (comprising its split OB3 and HJRL domains, Figure 1D, cartoons). Notably, this lobe is absent in many class averages, likely because it is connected to the RD domain by a flexible linker of 25 aa (Figure 1A) and may adopt many different positions relative to POT1, causing it to become averaged out in these classes (Figure 1—figure supplement 1A).

To further resolve elements within 2D-class averages of POT1/TPP1^N^, we incubated it with a synthetic 5’-biotin ssDNA and streptavidin to add a feature of distinct size and shape to the complex which should lie close to OB1 of POT1 (Lei et al., 2004). The streptavidin-ssDNA-POT1/TPP1^N^ complex eluted as three main peaks (Figure 1—figure supplement 2A): a small first peak consistent with a complex containing two POT1/TPP1^N^ complexes bound to a single streptavidin tetramer, followed by a second larger peak consistent with a complex containing one POT1/TPP1^N^ and one streptavidin tetramer, and finally another large peak containing excess streptavidin-ssDNA (Figure. 1E; Figure 1—figure supplement 2). Negative-stain EM analysis of a middle-peak fraction revealed class averages containing clear signal for a single streptavidin tetramer adjacent to an oblong bipartite lobe, from which another oblong lobe extends in many different directions (Figure 1E; Figure 1—figure supplement 1B). Based on the position of the biotin and known crystal structures, we interpret the bipartite oblong lobe to represent OB1 and OB2 of POT1 and the larger lobe to be its C-terminal domain bound to TPP1 (Figure 1E, cartoons). The OB fold of TPP1 is not visible in these averages, likely because the streptavidin-POT1 OB1/OB2 features dominate the alignment and TPP1 OB fold are averaged out. The 2D-class averages show complexes in a near-continuum of conformations from completely closed to fully extended as well as a conformation in which the C-terminal domain is bent backwards (Figure 1E; Figure 1—figure supplement 1B). These data are consistent with a high degree of flexibility of the hinge region between the N- and C-terminal domains of POT1, resulting in high conformational variability.

### Variable Association of the TIN2 TRFH Domain with POT1/TPP1

We performed similar analyses on POT1/TPP1/TIN2 (Figure 2A), hoping that we could exploit our domain assignments of POT1/TPP1^N^ to localize elements within the larger complex. Because removal of the GFP tag on TIN2 resulted in a poorly soluble complex, we purified the POT1/TPP1/GFP-TIN2 complex (Figure 2B,C), which was monodisperse in negative-stain EM images and gel filtration and specifically bound telomeric ssDNA (Figure 2—figure supplement 1). As with POT1/TPP1^N^, particles from this complex appeared homogenous in size and heterogenous in shape (Figure 2—figure supplement 1B). Since the ordered domains within the constituent proteins and the GFP tag are all similar in size and shape, we were unable to unambiguously assign identities to the features in the class averages from this complex (Figure 2—figure supplement 1B). To overcome this problem, we used the streptavidin-DNA labeling strategy as with POT1/TPP1^N^ (Figure 2D, top row; Figure 2—figure supplement 2A). As before, 2D-class averages showed streptavidin positioned near an oblong bipartite lobe, which we interpret as OB1 and OB2 of POT1. Certain classes were clearly relatable to 2D-class averages of POT1/TPP1^N^, with several resembling closed states of POT1/TPP1^N^ and others resembling more opened conformations (compare Figure 2D, top row, with Figure 1E). In all cases we observe additional signal, which could either represent GFP or the TRFH domain of TIN2.

**Figure 2.**
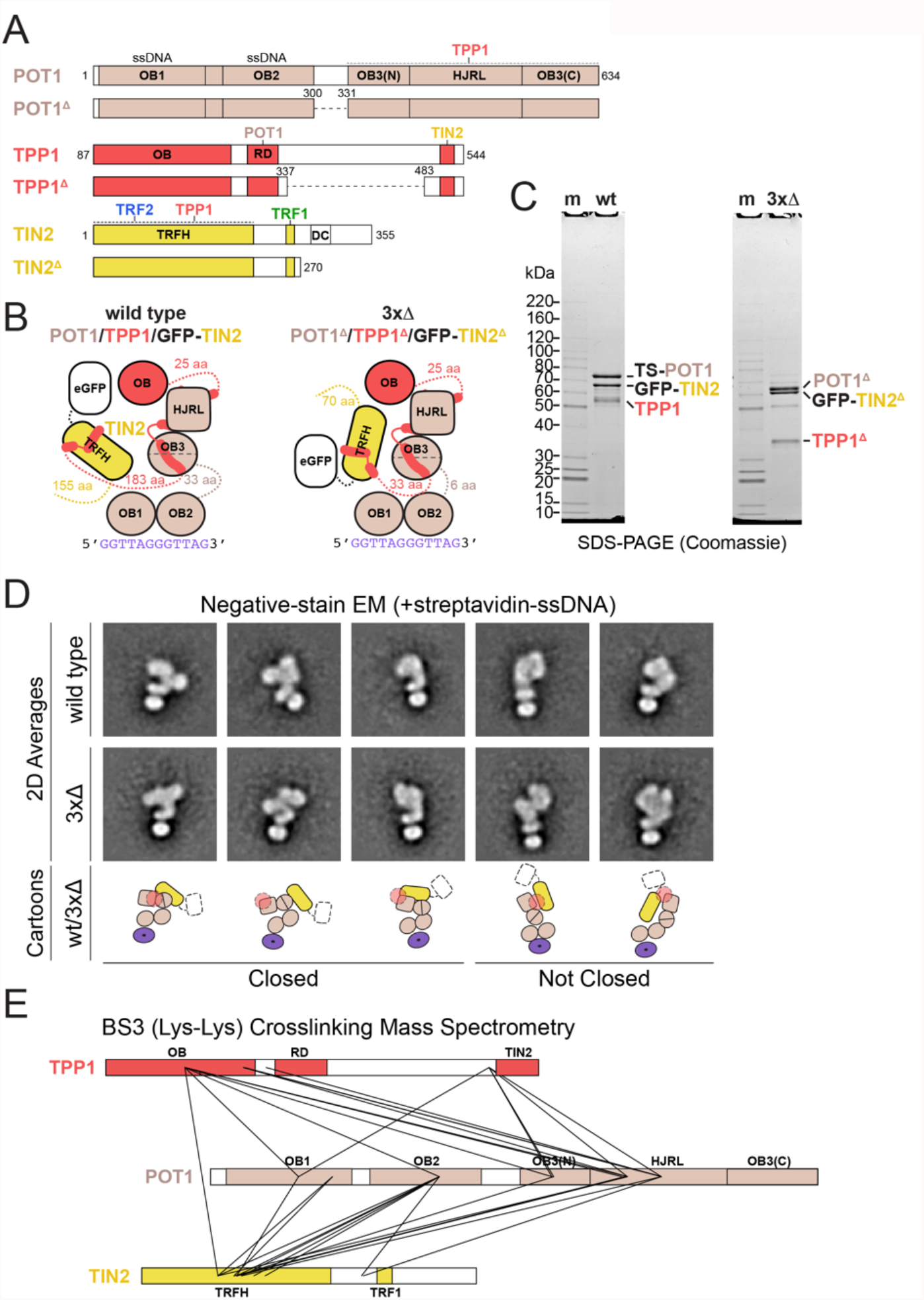
Structural variability of POT1/TPP1/GFP-TIN2. **(A)** Domain schematics for POT1, TPP1, TIN2, and deletion constructs with amino acids indicated. Coloring is as in panel 1A. In POT1 and TPP1 deletion constructs, dashed lines represent regions that have been replaced with (Gly-Gly-Ser)_2_. TRFH: TRF homology domain.**(B)** Cartoon schematics for POT1/TPP1/GFP-TIN2 wild type (wt) and deletion construct (3×Δ) bound to telomeric ssDNA. **(C)** SDS-PAGE of purified wt and 3×Δ POT1/TPP1/GFP-TIN2. TS: Twinstrep. Gel is 8-16% polyacrylamide Tris-glycine and stained with Coomassie blue. Molecular-weight markers are shown in kDa. **(D)** Negative-stain EM analysis of wt and 3×Δ POT1/TPP1/GFP-TIN2 bound to streptavidin-5’Biotin(GGTTAG)_2_ DNA. Reference-free 2D-class averages that were directly comparable to those for POT1/TPP1N were selected for display with cartoon interpretations below. **(E)** Crosslinking mass spectrometry of wt POT1/TPP1/GFP-TIN2 using the homo-bifuctional lysine-reactive reagent BS3. Domains are shown and colored as in panel A and crosslinks are shown as solid lines. Intramolecular crosslinks are not shown.

We asked whether the conformational heterogeneity of the POT1/TPP1/TIN2 complex was due to the flexible linkers acting as hinges in the three proteins. A POT1^Δ^/TPP1^Δ^/GFP-TIN2^Δ^ (referred to as 3×Δ) lacking the major disordered regions (POT1 residues 301-329, TPP1 residues 338-482, and TIN2 residues 271-355) (Figure 2A-C) was purified to apparent homogeneity (Figure 2C) and appeared as a monodisperse complex in negative-stain EM images (Figure 2—figure supplement 2B). Single-particle analysis using streptavidin-DNA labeling revealed that this complex also exhibits a wide-range of structural variability (Figure 2—figure supplement 2B), and that many reference-free 2D-class averages closely resemble those of the wild-type complex (Figure 2D, compare top and middle row). We conclude that POT1/TPP1/TIN2 remains conformationally variable in the absence of much of the flexible linker/hinge sequences.

We then subjected the POT1/TPP1/GFP-TIN2 complex to XLMS using the amine reactive molecule bis(sulfosuccinimidyl)suberate (BS3). We observed many intra- and inter-molecular crosslinks in the complex, including between TPP1 OB and POT1 HJRL as well as between TIN2 TRFH and the other components (Figure 2E; Supplemental Tables 2 and 3). Importantly, residues in TPP1 or TIN2 that are far apart in crystal structures crosslinked to the same or similar positions on POT1, indicating multiple conformations are present within the sample. For example, lysines 170 and 232 of TPP1 both crosslink to lysine 433 of POT1 (Figure 2E; Supplemental Table 3) despite being on opposite sides of the TPP1 OB fold (Wang et al., 2007). As an internal negative control, we do not observe any crosslinks between solvent-accessible lysines on the GFP or Twin-Strep-tags to the complex, confirming the specificity of the interactions we observe. The crosslinking data confirm the conclusion derived from the negative-stain EM analysis that the addition of TIN2 to POT1/TPP1 yields a complex with a high degree of conformational heterogeneity. These data also support the conclusion that the extra signal we see in class averages of POT1/TPP1/GFP-TIN2 relative to POT1/TPP1^N^ is the TRFH domain of TIN2 rather than GFP (Figure 2D, cartoons).

### POT1/TPP1/TIN2/TRF1 Bound to Telomeric DNA is a Dimeric Complex with High Conformational Heterogeneity

A complex containing POT1, TPP1, TIN2, and TRF1 (henceforth ‘the TRF1 complex’) was co-expressed in insect cells and purified to apparent homogeneity at high stringency (350 mM NaCl) (Figure 3B; Figure 3—figure supplement 1). To assess the compositional heterogeneity and stoichiometry of our preparation, we employed mass photometry (Young et al., 2018; Sonn-Segev et al., 2020). Purified POT1/TPP1/GFP-TIN2/TRF1 showed four peaks at measured masses of 80 kDa, 170 kDa, 270 kDa, and 470 kDa, which correspond to (TRF1)_2_ (97 kDa), POT1/TPP1/GFP-TIN2 (191 kDa), POT1/TPP1/GFP-TIN2/(TRF1)_2_ (288 kDa), and (POT1/TPP1/GFP-TIN2/TRF1)_2_ (480 kDa), respectively (Figure 3D; Supplementary Table 4). We performed native MS analysis and detected assemblies with these stoichiometries at high mass accuracy and resolution (Figure 3E; Figure 3—figure supplement 1C; Supplementary Table 5). Overall, we observed that the TRF1 dimer can associate with one or two POT1/TPP1/GFP-TIN2 assemblies and that the TRF1 complex partially dissociates during purification.

**Figure 3.**
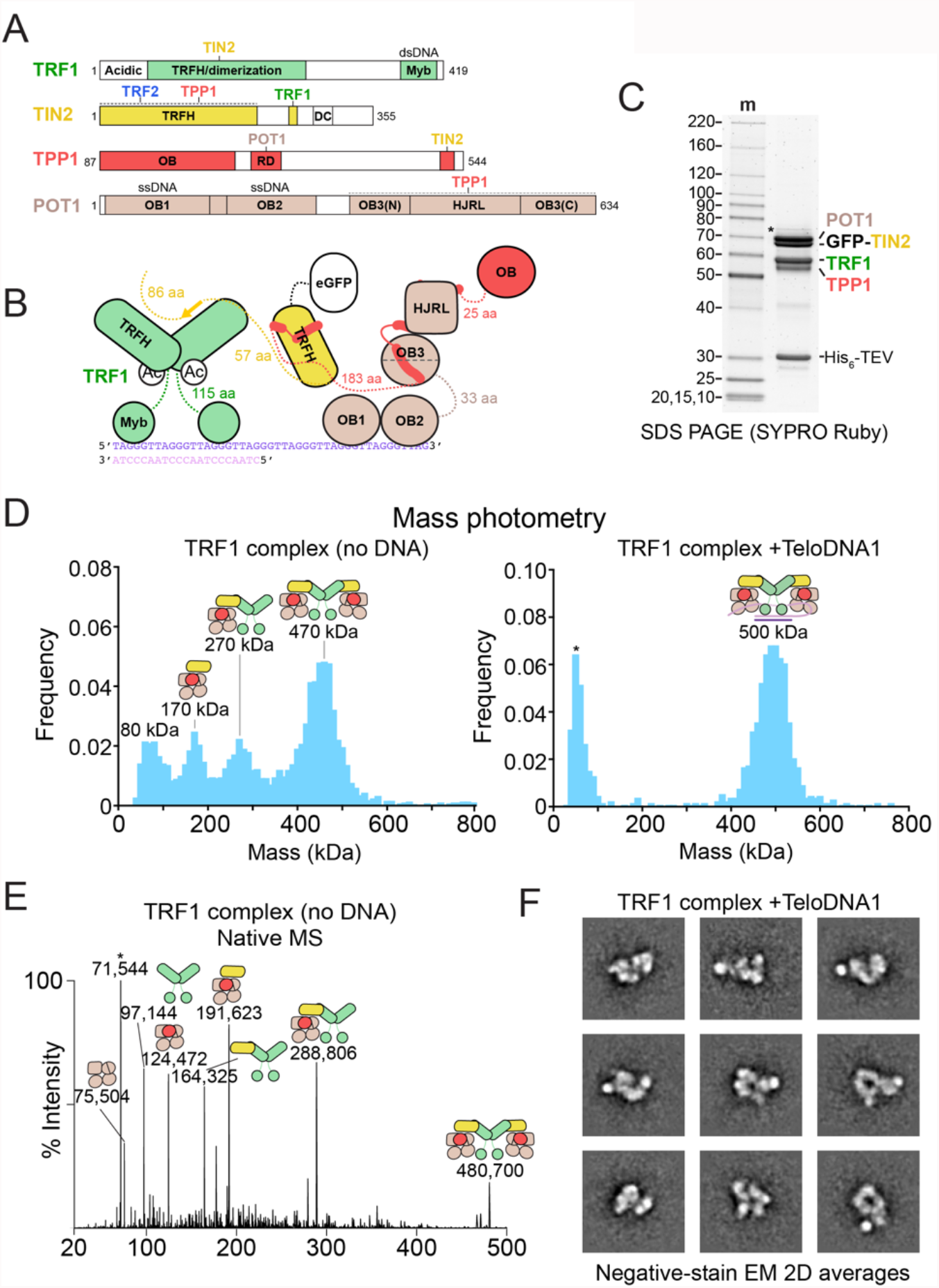
Structural heterogeneity within the TRF1 complex. **(A)** Domain schematics for POT1, TPP1, TIN2, and TRF1 with amino acids indicated. **(B)** Cartoon schematic for POT1/TPP1/GFP-TIN2/TRF1 bound to a ds-ssDNA junction **(C)** Purified POT1/TPP1/GFP-TIN2/TRF1 prior to glycerol-gradient purification fractionated on SDS-PAGE. Gel is 8-16% polyacrylamide Tris-glycine and stained with SYPRO Ruby. Molecular-weight markers are shown in kDa. **(D)** Mass photometry of TRF1 complex purified in the presence or absence of ds-ss junction telomeric DNA (TeloDNA1). Final protein concentration was 12 nM (by GFP absorbance) and peak maxima are indicated along with cartoons that correspond to complexes of that molecular weight to within ~10%. **(E)** Native MS analysis of reconstituted TS-POT1/TPP1/GFP-TIN2/TRF1. The measured masses (in Da) are indicated along with cartoons that correspond to complexes of that approximate molecular weight to within 0.1% (see Supplementary Table 5). Asterisk indicates a species that likely corresponds to HSP70 contamination. **(F)** Negative-stain EM analysis of POT1/TPP1/GFP-TIN2/TRF1 bound to TeloDNA1. Reference-free 2D-class averages showing a high signal-to-noise ratio were selected for display.

We then sought to stabilize the dimeric complex by adding a DNA containing both ds- and ssDNA of telomeric sequence. We found that DNA containing an optimal TRF1-binding site (Bianchi et al., 1999) adjacent to a (GGTTAG)_)4_ ssDNA 3’ overhang gave rise to a single species when bound to the TRF1 complex in a native gel (hereafter ‘TeloDNA1’, Figure 3— figure supplement 1B). When TeloDNA1 was added prior to ultracentrifugation, the purified TRF1 complex behaved as a single 500-kDa species in mass photometry (Figure 3D), which is close to the predicted molecular weight of (POT1/TPP1/GFP-TIN2/TRF1)_2_ bound to TeloDNA1 (505 kDa). Similarly, addition of TeloDNA1 greatly increased the formation of the 500-kDa complex when the TRF1 complex was reconstituted by mixing purified TRF1 with POT1/TPP1/GFP-TIN2 (Figure 3—figure supplement 1D).

The TRF1 complex bound to TeloDNA1 was purified by glycerol-gradient ultracentrifugation for analysis by negative-stain EM (Figure 3—figure supplement 1E; Figure 3—figure supplement 2A). Peak fractions contained all four protein components and a native gel stained for nucleic acid demonstrated the presence of a single species bound to DNA (Figure 3—figure supplement 1E). Negative-stain EM images revealed that the particles were homogenous in size and did not aggregate, but 2D-class averages showed extensive heterogeneity (Figure 3F; Figure 3—figure supplement 2A). At 200 mM KCl in the absence of DNA, the TRF1 complex tended to aggregate (Figure 3—figure supplement 1E; Figure 3—figure supplement 2B), and 2D-class averages of the non-aggregated particles displayed less defined features compared to the DNA-bound complex (Figure 3—figure supplement 2B).

For streptavidin-DNA labeling of the TRF1 complex, two POT1-binding sites were included in the DNA construct so that the two POT1 proteins in the dimeric complex could engage DNA substrate. Negative-stain EM of POT1/TPP1^N^ bound to this substrate showed streptavidin close to two oblong bipartite lobes, as expected from the binding of pairs of POT1 OB1 and OB2 (Figure 3—figure supplement 3A). In some cases, the two sets of OB folds adopted a curved U-or V-shape consistent with the DNA path observed in the crystal structure of the OB folds bound to DNA (Lei et al., 2004). Using this feature as a guide, we analyzed POT1/TPP1/GFP-TIN2 bound to this substrate (Figure 3—figure supplement 3B). Class averages of this sample showed larger complexes than those for POT1/TPP1^N^, and while many classes contained streptavidin associated with a U-feature, other elements were poorly resolved. Adding TRF1 to the reconstitution further decreased the quality of alignment within 2D classes, indicating that streptavidin-DNA labeling using this substrate is not a suitable strategy to identify elements within 2D-class averages of the TRF1 complex (Figure 3—figure supplement 3C).

### Conformational heterogeneity and stoichiometry of shelterin

The full shelterin complex (Figure 4A-C) was reconstituted by mixing purified TRF2/Rap1 (Figure 4—figure supplement 1A) with the TRF1 complex in the presence and absence of a telomeric DNA containing four Myb-binding sites and two POT1-binding sites (TeloDNA2) (Figure 4—figure supplement 1; Figure 4—figure supplement 2). Given that the TRF2-TIN2 interaction is disrupted at high salt (Ye et al., 2004a), the reconstituted complexes were diluted in buffer containing 150 mM KCl and this salt concentration was maintained in the glycerol gradient.

**Figure 4.**
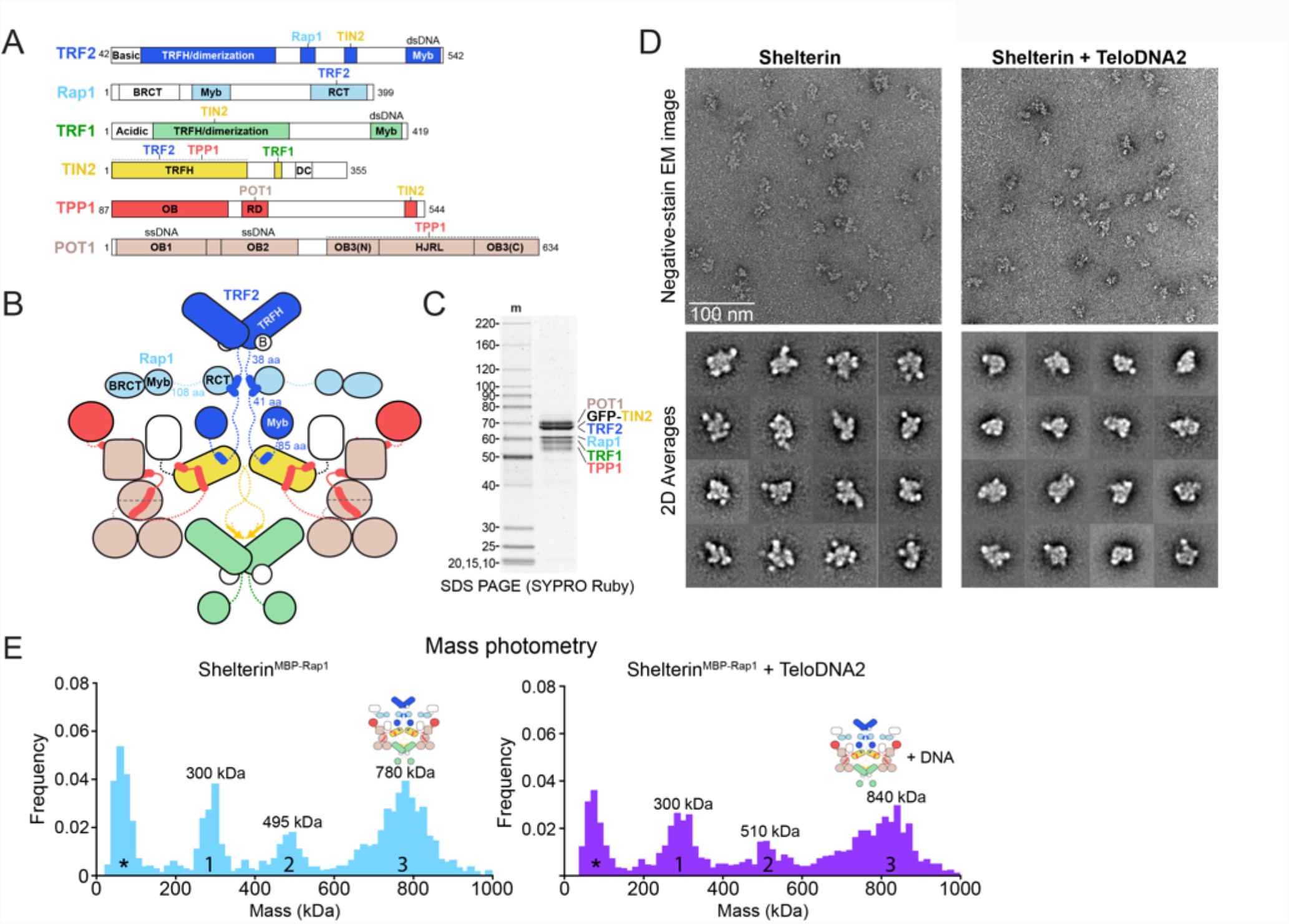
Structural heterogeneity within shelterin. **(A)** Domain schematics for TRF2, Rap1, TRF1, TIN2, TPP1, and POT1 with amino acids indicated. RCT: Rap1 C-terminal domain; BRCT: BRCA1 C-terminal domain. **(B)** Cartoon schematic for dimeric shelterin. Coloring is as in panel 1A. **(C)** SDS-PAGE of purified reconstituted shelterin. Gel is 8-16% polyacrylamide Tris-glycine and stained with SYPRO Ruby. Molecular-weight markers are in kDa. **(D)** Negative-stain EM analysis of shelterin purified in the presence or absence of ds-ss junction telomeric DNA (TeloDNA2). EM images displaying raw particles are shown above reference-free 2D-class averages. Class averages showing a high signal-to-noise ratio were selected for display. **(E)** Mass photometry of reconstituted shelterin containing MBP-Rap1 and GFP-TIN2 after an amylose pulldown. Data were taken in the presence or absence of exogenously added ds-ss junction telomeric DNA (TeloDNA2) as indicated. Peaks are numbered and approximate maxima are indicated. Asterisks indicate a peak arising from minor buffer contaminants. Zinder et al., Structural Heterogeneity of Human Shelterin

In the absence of DNA, fractions containing TRF2 and Rap1 were observed co-migrating with components of the TRF1 complex at much higher glycerol concentrations than for TRF2/Rap1 alone (Figure 4—figure supplement 1A). Unlike the TRF1 complex, DNA-free shelterin was both soluble and monodisperse at low salt. However, negative-stain EM analysis of a peak fraction from the glycerol gradient showed that shelterin behaved as a heterogenous complex in 2D-class averages (Figure 4D; Figure 4—figure supplement 1).

TeloDNA2-bound shelterin was more compact than the apo complex and migrated to higher density, but it remained extensively heterogenous in 2D-class averages (Figure 4D; Figure 4—figure supplement 2). Thus, the addition of the TRF2/Rap1 components and/or DNA to the TRF1 complex prevented aggregation at physiological salt concentration but did not significantly limit conformational variability.

We then sought to probe the stoichiometry of the full complex, but the complex quantitatively bound to membranes during concentration/buffer exchange by spin filtration. To overcome this issue, we reconstituted shelterin by mixing the TRF1 complex with TRF2/His_6_MBP-Rap1, loading the sample on a glycerol gradient and performing an amylose pulldown on the peak fractions (Figure 4—figure supplement 3A,B). The isolated complex was analyzed by mass photometry, which showed three prominent peaks at 300, 495 and 780 kDa (not counting an additional low molecular weight peak from a buffer contamination) (Figure 4E; Figure 4—figure supplement 3C). Peak 1 would be consistent with the molecular masses of either (TRF2/His_6_MBP-Rap1)_2_ (286 kDa) or POT11/TPP1/GFP-TIN2/(TRF1)_2_ (285 kDa), peak 2 with either POT11/TPP1/GFP-TIN2/(TRF2/His_6_MBP-Rap1)_2_ (474 kDa) or (POT1/TPP1/GFP-TIN2/TRF1)_2_ (473 kDa), and peak 3 with (POT1/TPP1/GFP-TIN2/TRF1/TRF2/ His_6_MBP-Rap1)_2_ (759 kDa) (Supplementary Table 4). Addition of TeloDNA2 to this complex resulted in shifts of peaks 2 and 3 of approximately the molecular weight of the DNA (41 kDa) further confirming our peak assignments (Figure 4E, Figure 4—figure supplement 3C). From these observations we conclude that shelterin can assemble into a fully dimeric complex.

## Discussion

This study reports on the structure of shelterin and its subcomplexes reconstituted *in vitro*. No DNA was required to form the six-subunit shelterin complex, further confirming that shelterin can self-assemble in the absence of its telomeric DNA substrate (Erdel et al., 2017). Single-particle negative-stain EM and XLMS data revealed that shelterin and its subcomplexes can adopt a wide array of conformations that are not altered by the addition of optimal DNA substrates. Due to this extreme structural heterogeneity (Figure 5), high-resolution structure determination of shelterin is currently intractable. Nonetheless, the data reported here provide important insights into the architecture and functions of shelterin.

**Figure 5.**
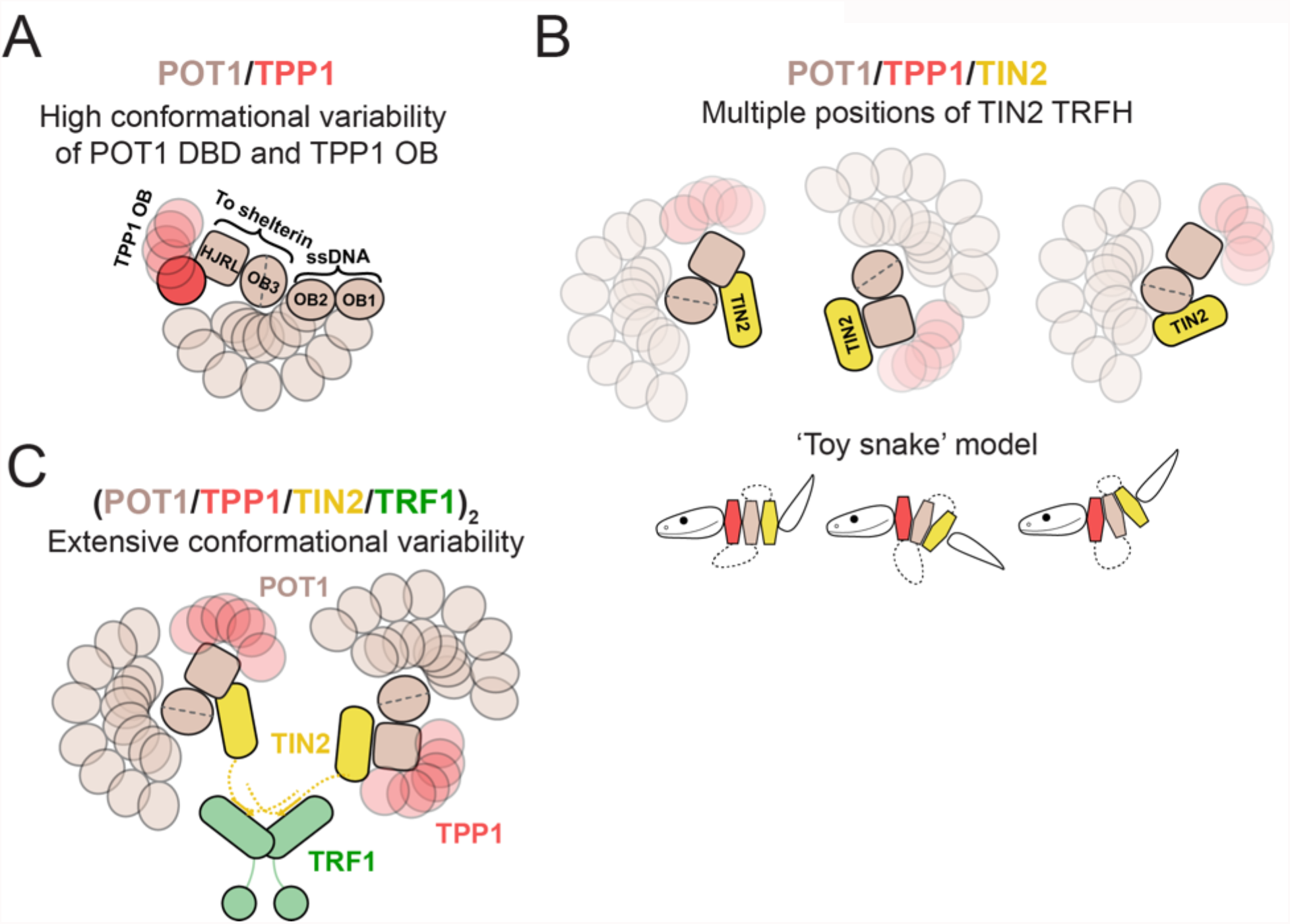
Models of structural heterogeneity in human shelterin. **(A)** Visual depiction of conformational variability within POT1/TPP1^N^. In these models the C-terminal domains of POT1 are arbitrarily used as an anchor. Conformational variability of the DNA-binding OB folds and the TPP1 OB fold is depicted as an ensemble of possible conformations that these features can adopt relative to that anchor. **(B)** Models for flexibility within POT1/TPP1/TIN2. The TRFH domain of TIN2 is shown occupying different discrete positions on the POT1 anchor. For the ‘toy snake’ model, structured domains are depicted as hexagonal snake segments with flexible linkers shown as dotted lines connecting them. These domains can associate with each other in a variety of conformations, and truncation of flexible linkers would result in only minor changes in structure and conformational variability. **(C)** Model for structural heterogeneity of the TRF1 complex. The POT1/TPP1/TIN2 subcomplex remains flexible and TIN2 is presumably flexibly tethered to TRF1.

The heterodimer formed by full-length POT1 and the N-terminal half of TPP1 (comprising its OB fold and RD) showed a large array of distinct conformations. The 2D-class averages suggest a continuum of different positions of the C-terminal half of POT1 (composed of the split OB3/HJRL domain to which the TPP1 RD binds) and the two N-terminal OB folds that bind to ss telomeric DNA (Figure 5A). Clearly, the association of TPP1 with the split OB3/HJRL domain of POT1 does not lead to a rigid POT1 structure. Therefore, POT1 may be best viewed as a DNA-binding module (OB1 and OB2) that is flexibly tethered to shelterin via the interaction of its split OB3/HJRL domain with TPP1. This architecture could enable POT1 to bind to ssDNA and fulfill its functions with greater versatility than if it were rigidly attached and need to be specially positioned by the dsDNA binding components.

Reconstitution of the POT1/TPP1/TIN2 complex with full-length proteins resulted in a trimer with 1:1:1 stoichiometry, as expected. The structure of the POT1/TPP1 heterodimer was readily discerned within the heterotrimer, indicating that TIN2 does not significantly affect the positional variation of the POT1 halves. Interestingly, negative-stain EM and XLMS data revealed that the TRFH domain of TIN2 (which binds to the C-terminal region of TPP1) is found at many different positions in close association with POT1 (Figure 5B). The TIN2-binding site in TPP1 is separated from the POT1-binding RD domain by an unstructured and poorly conserved region of 150 aa. However, TIN2 is not freely diffusing around POT1/TPP1 on this tether but has several preferred positions yielding well-defined negative-stain EM class averages and high-confidence crosslinks between the TIN2 TRFH domain and all domains of known structure in POT1/TPP1. Consistent with the view that TIN2 is not randomly moving around POT1/TPP1, deletion of the 150-aa region of TPP1 did not have a significant effect on the negative-stain EM class averages. These data suggest that POT1/TPP1/TIN2 may contain a core (comprised of POT1 OB3/HJRL, TPP1 OB+RD, and TIN2 TRFH) which, while variable in structure, may have a limited range of conformations. Contrasting with a ‘beads-on-a-string model’, the behavior of the POT1/TPP1/TIN2 core resembles that of a toy wooden snake (Figure 5B), in which the segments would represent structured domains that associate closely but can adopt many different conformations relative to each other. In this analogy, the flexible regions tethering the structured modules would be looping out from them such that their removal would leave the structure and flexibility of the complex unchanged. Future studies using time-resolved measurements on conformational states will be needed to determine if individual complexes can toggle freely between conformational states.

Addition of TRF1 to the POT1/TPP1/TIN2 complex yielded a shelterin subcomplex that could have a fully dimeric stoichiometry (Figure 5C). We also observed complexes containing only one POT1/TPP1/TIN2 trimer bound to the TRF1 homodimer, consistent with observations of the TRF2-containing subcomplex (Lim et al., 2017). However, in both the apo state and when bound to a DNA substrate containing a TRF1- and two POT1-binding sites, the dimeric complex dominated, which was also the case for the full shelterin complex. Because shelterin components are expressed at different levels in cells (Takai et al., 2010) and ssDNA is relatively scarce at the telomere, it is likely that it can adopt multiple stoichiometries. This is further supported by the observation that breaking of the TIN2 bridge at either the TRF1 or TRF2 site results in minor or no problems with telomere protection and maintenance, so long as POT1/TPP1/TIN2 can efficiently be recruited to the telomere (Frescas & de Lange, 2014; Takai et al., 2011). Structurally, addition of TRF1 did not significantly decrease the variability of the POT1/TPP1/TIN2 complex, resulting in many different negative-stain EM class averages with features that we were unable to unambiguously assign. Similarly, reconstitution of the full shelterin complex by addition of TRF2/Rap1 resulted in a great degree of structural variability regardless of whether it was bound to a DNA substrate.

A limitation of this study is that it assumes that there are no missing components of shelterin, which could, theoretically, lend conformational homogeneity to the complex. From numerous proteomic studies it is clear that there are no telomere-specific factors missing. However, it is not excluded that proteins that are not specific to telomeres play a role in the structure of the complex. Recent examples of this phenomenon are the telomerase and *γ*-TURC complexes, each of which were purified from human cells and co-purified with abundant endogenous components that enhanced their stability (H2A/H2B for telomerase and actin for *γ*-TURC) (Ghanim et al., 2021; Wan et al., 2021; Wieczorek et al., 2020; Liu et al., 2020). Additionally, it is possible that elements of shelterin are stabilized in discrete conformations when bound to one or more of its many accessory factors or when in association with the telomeric DNA maintenance factors (telomerase and CST/Pola/primase).

All known interactions within shelterin occur between small domains or peptides flanked by disordered regions and structured domains, and not between two structured domains (de Lange, 2018; Lim & Cech, 2021; Hu et al., 2017). This study uncovered an association between the TRFH domain of TIN2 with the structured domains of POT1/TPP1, but this interaction was found to be highly variable (Figure 5B, toy snake model). These features of shelterin likely underlie the observation that increasingly larger shelterin complexes resulted in increasing structural variability, which decreased the quality of alignment in single-particle analysis and precluded localization of structured domains within the larger complexes.

Why is shelterin so flexible? We speculate that structural heterogeneity is a beneficial feature of the complex that permits shelterin to fulfill its many functions. For example, flexibility may enable shelterin to accommodate the many conformations adopted by telomeric DNA. Shelterin binds to telomeres in their linear state, associating with ds repeats that are largely chromatinized and with the ds-ss transition and 3’ overhang at the terminus. Shelterin-bound telomeres also can adopt a t-loop configuration, featuring a D-loop as well as branched DNA at the t-loop base. Furthermore, the telomeric long non-coding RNA, TERRA, can form R-loops in telomeres, which may present additional challenges to shelterin (Azzalin et al., 2007; Azzalin & Lingner, 2015). Finally, shelterin needs to accommodate numerous interacting partners involved in the maintenance of telomeric DNA (e.g., telomerase, CST, Apollo, BLM, RTEL1), which may require different conformations to position each on their respective DNA substrates.

## Materials and Methods

### Cloning and protein expression

All constructs were cloned for baculovirus-based expression of multicomponent complexes in insect cells using the biGBac system (Weissmann et al., 2016). Briefly, DNA fragments encoding proteins, tags, and/or protease-cleavage sites were generated by PCR and cloned between a polyhedrin promoter and SV40 terminator in a subcloning vector (pLIB) using Gibson assembly (Gibson et al., 2009). Gene-expression cassettes containing the promoter, expression construct, and terminator were amplified from these vectors by PCR using a predefined primer set to allow Gibson assembly into another vector (pBIG1). In the case of POT1/TPP1/GFP-TIN2/TRF1, these vectors were digested with PmeI and further assembled into a final expression vector (pBIG2).

All insect cell incubation steps were performed at 27°C and high humidity (68-74%), and all transfections/infections were performed on cells in logarithmic growth. Vectors were converted into purified bacmids using DH10Bac (Thermo Fisher) cells per manufacturer’s instructions. Bacmids were transfected into adherent Sf9 cells grown in Sf-900 II serum-free medium (Thermo Fisher) using CellFectin II (Thermo Fisher). Four days post transfection, the medium was harvested, centrifuged, filtered through 0.45 µm, aliquoted and stored at 4°C in the dark for up to 12 months (P1 Virus). 1.5 mL of P1 stock was then used to infect 30e6 adherent Sf9 cells in Sf-900 II medium. Three days after infection with P1, the medium was harvested, centrifuged, filtered through 0.45 µm, aliquoted and stored at 4°C in the dark for up to 6 months (P2 Virus). 3 mL of P2 stock was then used to infect 60e6 adherent Sf9 cells in Sf-900 II medium. Three days after infection, the medium on top of the adherent Sf9 cells was harvested, centrifuged (P3 virus), and added to 400 mL Tni cells at 2.0 cells/mL in ESF900 medium and grown with spinning at 150 RPM. Tni cells were harvested 72 hrs post infection by centrifugation, transferred to a blocked syringe, then transferred dropwise into storage containers submerged in liquid nitrogen and stored at −80°C until use. Cells were verified for expression by visual inspection for fluorescent constructs or Western blotting of whole-cell lysates for others.

### Protein purification

Prior to purification, frozen cells were disrupted by cryogenic milling (Retsch) and approximately half of the resulting powder of a 400-mL culture was used for purification while the other half was stored at −80°C. All protein-purification steps were performed on ice or at 8°C and using chilled buffers. All protein concentrations were determined by A_280_ and a calculated extinction coefficient from https://web.expasy.org/protparam/ for complexes lacking GFP or A_488_ for GFP-containing complexes using an extinction coefficient for eGFP at 488 nm of 53,000 M^−1^cm^−1^. UV-vis absorbance measurements were made using a NanoDrop ND-1000 (Thermo Fisher). All buffers used in FPLC were degassed using a benchtop vacuum and cooled to 8°C prior to use. Glycerol gradients were mixed using a Gradient Master (BIOCOMP) at room temperature and cooled to 4°C prior to use. All polyacrylamide gels were purchased from Thermo Fisher, and Benchmark unstained protein ladder (Thermo Fisher) was used as a molecular weight marker for SDS-PAGE. After the final purification step, samples were concentrated by centrifugal filtration, aliquoted, flash frozen, and stored at −80°C.

POT1/TPP1^N^ was purified by GFP affinity pulldown (resin made in-house) followed tag cleavage with TEV protease (purified in-house) and gel filtration (Cytiva). POT1/TPP1/GFP-TIN2 was purified by streptactin affinity pulldown (IBA biosciences), heparin (Cytiva), and gel filtration. POT1/TPP1/GFP-TIN2 3×Δ was purified by streptactin affinity pulldown followed by strong-anion exchange (Cytiva), TEV cleavage, and gel filtration. POT1/TPP1/GFP-TIN2/TRF1 was purified by streptactin affinity pulldown, mCherry affinity pulldown (resin made in-house), tag cleavage with TEV protease, and ultracentrifugation through a 10-30% glycerol gradient. TRF1 and TRF2/Rap1 were purified by amylose pulldown (New England BioLabs), heparin, tag cleavage with 3C protease (purified in-house), and gel filtration. TRF2/His_6_MBP-Rap1 was purified by nickel-NTA (Qiagen), heparin, and gel filtration.

### DNA substrates

All unlabeled oligonucleotides were purchased from Thermo Fisher and purified using standard desalting unless otherwise specified. All 5’Biotin oligonucleotides were ordered from Thermo Fisher with HPLC purification. All single-stranded 5’ Cy5.5 DNAs were ordered from Integrated DNA Technologies with HPLC purification. The Cy5.5-labeled telomeric dsDNA was generated by mixing 540 pmol of HPLC purified 5’CATCAATAGGGTTCATCCTAGGGTTGTACTG3’ DNA with 650 pmol of HPLC purified 5’CAGTACAACCCTAGGATGAACCCTATT3’ in 25 µL annealing buffer (10 mM HEPES-KOH pH 7.5, 100 mM NaCl, 0.1 mM EDTA, 0.2 µm filtered), heating to 98°C for 2 min, and cooling quickly to 12°C in a thermocycler. The annealed product was labeled by fill-in synthesis using 5 U Klenow exo-(New England Biolabs) and a dNTP mix containing 40 µM each of dATP, dCTP, dGTP and SulfoCy5.5-dUTP (Lumiprobe). These reactions were then exchanged into annealing buffer using a BioSpin P6 column (Bio-Rad) and the concentration was attained by measuring A_673_ and using an extinction coefficient of 211,000 M^−1^cm^−1^. To make TeloDNA1, 7 nmol of 5’CATCAATAGGGTTCATCACTAGGGTTAGGGTTAGGGTTAGGGTTAGGGTTAG3’ was annealed to 10.5 nmol of 5’CTAACCCTAGTGATGAACCCTATTGATG3’ by heating to 98°C and cooling quickly to 12°C in 100 µL annealing buffer. The volume was then adjusted to 550 µL with annealing buffer and injected onto a Superdex200 10/300 GL column equilibrated in phosphate-buffered saline (PBS, 4.3 mM Na_2_HPO_4_, 1.5 mM KH_2_PO_4_, 140 mM NaCl, 2.7 mM KCl, pH 7.4). Fractions were analyzed by TBE-PAGE with SYBR Gold (Invitrogen) staining and those containing primarily the annealed product were pooled, concentrated to 100 µL in a Microcon YM-10, and stored at −20°C. TeloDNA2 was generated in the same way, except the sequences used were 5’CCTATCTAGGGTTTTCTACTAGGGTTCATCAATAGGGTTCATCACTAGGGTTAGGGTTAGGGTTAGGGTTAGGGTTAG3’ and 5’CTAACCCTAGTGATGAACCCTATTGATGAACCCTAGTAGAAAACCCTAGATAGG3’.

### Electrophoretic mobility shift assays (EMSAs)

For POT1/TPP1^N^ and POT1/TPP1/GFP-TIN2, variable concentrations of protein were mixed with 0.25 nM Cy5.5-labeled DNA in binding buffer (20 mM HEPES-KOH pH 7.5, 100 mM KCl, 0.5 mM MgCl_2_, 0.5 mM TCEP-HCl, 0.05% v/v IGEPAL Co-630, 8% glycerol, 50 µg/mL bovine serum albumin (New England BioLabs)) and incubated for 30 min at room temperature (~22°C). For POT1/TPP1^N^, 2.5 µL was separated on a 4-20% polyacrylamide-TBE gel in cold 100 mM Tris-Borate pH 8.3. For POT1/TPP1/GFP-TIN2, 6 µL was separated on 0.6% agarose-Tris-Borate in cold 100 mM Tris-Borate pH 8.3. Gels were imaged using the near-IR setting of a Typhoon scanner (GE). For the qualitative EMSA of POT1/TPP1/GFP-TIN2/TRF1, variable concentrations of protein were mixed with 50 nM of the indicated DNA in binding buffer containing 200 mM KCl and incubated on ice for 30 min. 5 µL of each protein/DNA complex was then separated on 4-20% polyacrylamide TBE gels run in agarose-TBE in cold Tris-Borate buffer, the gel stained with SYBR gold per manufacturer’s instructions, and imaged using the ethidium bromide setting of an Alpha imager (Alpha Innotech). For native gels of glycerol gradient fractions, 5 µL of fractions were run and imaged as with the qualitative TRF1 complex EMSA.

### Sample preparation for negative-stain EM

For POT1/TPP1^N^, an aliquot of purified protein was thawed, centrifuged at 20,000 x g, and the concentration of the supernatant was measured. 550 pmol of protein was then added to a tube containing 1.1 nmol of 5’Biotin(GGTTAG)_2_ DNA (Thermo Fisher) in a volume of 100 µL dilution buffer (20 mM HEPES-KOH pH 7.5, 200 mM KCl, 0.1 mM TCEP-HCl, 10% glycerol). This mixture was then incubated on ice for 10 min, after which point the volume was adjusted to 550 µL with dilution buffer (final concentrations: 1 µM protein complex, 2 µM DNA). After centrifugation at 20,000 x g for 1 min, the supernatant was injected onto a Superose 6 increase 10/300 GL column (Cytiva) that had been equilibrated in degassed dilution buffer. Fractions were collected and analyzed by SDS-PAGE and those of interest were analyzed by negative-stain EM. If necessary, samples were diluted with dilution buffer to a concentration that achieved a high density of well-separated particles on the grid. For streptavidin labeling, 1.1 nmol of DNA was mixed with 3.3 nmol of D-biotin-Tris pH 7.5, and then 2.2 nmol purified streptavidin (New England BioLabs) was added. This mixture was incubated on ice for 10 min, then another 10 nmol of biotin added and the volume adjusted to 100 µL. Subsequent steps were performed as before. When 5’Biotin(GGTTAG)_4_ DNA was used, the concentrations of DNA, D-biotin, and streptavidin were reduced by a factor of four. For POT1/TPP1/GFP-TIN2 wt and 3×Δ, a similar protocol was used, except that final concentrations were 0.4 µM complex, 0.8 µM 5’Biotin(GGTTAG)_2_ DNA, 1.6 µM streptavidin (if present), and 2.4 µM D-biotin (if present).

For POT1/TPP1/GFP-TIN2/TRF1, the eluate from the mCherry column after concentration was used as an input. 220 pmol of the eluate (by GFP absorbance at 488 nm) was added to 220 pmol TeloDNA1 or buffer and the volume adjusted to 220 µL with dilution buffer lacking glycerol (final concentrations are 1 µM GFP-TIN2 in the TRF1 complex by A_488_, 0 or 1 µM TeloDNA1). This mixture was incubated on ice for 2 hrs and then 200 µL was added to the top of 10-30% glycerol gradients in 20 mM HEPES-KOH pH 7.5, 200 mM KCl, 0.5 mM TCEP-HCl. The gradients were centrifuged and fractionated as before. Fractions were analyzed by SDS-PAGE with SYPRO Ruby staining and the DNA-containing sample was additionally analyzed by native TBE-PAGE with SYBR Gold staining. Fractions of interest were further analyzed by negative-stain EM.

For POT1/TPP1/GFP-TIN2 and POT1/TPP1/GFP-TIN2/TRF1 bound to streptavidin-5’Biotin(GGTTAG)_4_ DNA, 165 pmol of DNA was mixed with 500 pmol D-biotin, and then 330 pmol of streptavidin was added. The reaction was incubated for 10 min on ice and then another 1 nmol of D-biotin was added and the volume adjusted to 100 µL with dilution buffer. To this sample, 330 pmol of POT1/TPP1/GFP-TIN2 was added and the mixture incubated for 10 min on ice. Then, either buffer or buffer plus 165 pmol of (TRF1)_2_ was added to a final volume of 220 µL and the mixture was incubated on ice for another 1 hr before separation by ultracentrifugation, fractionation, and analysis as above.

For the shelterin reconstitution, the eluate from the mCherry step after concentration of the TRF1 complex and purified TRF2/Rap1 were used as inputs. 260 pmol the TRF1 complex was mixed with 130 pmol of (TRF2/Rap1)_2_ and incubated on ice for 10 min. Then, either buffer or buffer plus 200 pmol TeloDNA2 was added to a final volume of 220 µL (final concentrations were 1.2 µM GFP-TIN2 in the TRF1 complex by A488, 0.6 µM (TRF2/Rap1)_2_, 0 or 0.9 µM TeloDNA2). This sample was incubated on ice for 2 hrs then added to the top of 10-30% glycerol gradients in 20 mM HEPES-KOH pH 7.5, 150 mM KCl, 0.5 mM TCEP-HCl and separation by ultracentrifugation, fractionation, and analysis as above.

### Negative-stain EM data collection and image processing

Protein samples (3.5 µL) were adsorbed to glow-discharged carbon-coated copper/collodion grids, washed three times with water and once with freshly prepared 0.7% w/v uranyl formate, then stained for 30 s with 0.7% uranyl formate. Samples were imaged at room temperature using a Phillips CM10 electron microscope equipped with a tungsten filament and operated at an acceleration voltage of 80 kV. Images were collected with an AMT ActiveVu CCD camera at a calibrated pixel size of 2.8 Å. Particles were auto-picked using the swarm (for POT1/TPP1^N^ and POT1/TPP1/TIN2 wt/3×Δ) or Gauss (for other complexes) picker in EMAN2.1 (Tang et al., 2007). Particles were then extracted and subjected to 2-3 rounds of 2D classification in *RELION* 3.0 (Zivanov et al., 2018) to remove junk particles. After the final round of classification, particles were re-extracted using the updated coordinates from 2D classification in RELION, and that stack was used as input for 2D classification in ISAC 2.3.2 (Yang et al., 2012) using a pixel error threshold of 0.7 and variable minimum and maximum particle number per class (Supplemental Table 1). Classes containing readily interpretable features from the ISAC output were selected for display in main figures while all classes are displayed in the supplement.

### Crosslinking mass spectrometry

BS3 cross-linker (Proteochem, c1103) was dissolved in LC-MS grade H_2_O (Proteochem, LC6330) at 50 mM and added to POT1/TPP1/GFP prepared at 1 mg/mL in NHS-ester non-reactive buffer to the final concentration of 0.35-0.75 mM. Reactions were performed at 25°C in disposable inert cuvettes (UVette, Eppendorf), and monitored by continuous looped dynamic light scattering measurements of polydispersity (Pd<10 %; DynaPro NanoStar, Wyatt) (Meyer et al., 2015). Cross-linking was quenched after 30 min incubation by addition of Tris-HCl pH 8.0 to a final concentration of 5 mM.

Samples were dialyzed against 100 mM ammonium bicarbonate, reduced with 50 mM TCEP at 60°C for 10 min and alkylated with 50 mM iodoacetamide in the dark for 15 min at 37°C. Digestion was carried out at 37°C overnight with 125 ng/mL sequencing grade modified trypsin (Themo Fisher) in 25 mM ammonium bicarbonate supplemented with ProteaseMax (Themo Fisher). Reaction mix was supplemented with trifluoroacetic acid (TFA, Themo Fisher) to a final concentration of 0.1%. The resulting peptides were passed through C18 Spin Tips (Themo Fisher) before elution with 40 μL of 80% acetonitrile (ACN, Themo Fisher) in 0.1% TFA. Eluted peptides were dried and resuspended in 20 μL 0.1% formic acid (Themo Fisher) for MS analysis. Peptides were analyzed in an Orbitrap Fusion Lumos mass spectrometer (Themo Fisher) coupled to an EASY-nLC (Themo Fisher) liquid chromatography system, with a 2 μm, 500 mm EASY-Spray column. The peptides were eluted over a 120-min linear gradient from 96% buffer A (water) to 40% buffer B (ACN), then continued to 98% buffer B over 20 min with a flow rate of 250 nL/min. Each full MS scan (R = 60,000) was followed by 20 data-dependent MS2 scans (R = 15,000) with high-energy collisional dissociation (HCD) and an isolation window of 2.0 m/z. Normalized collision energy was set to 35. Precursors of charge state ≤ 3 were collected for MS2 scans in enumerative mode, precursors of charge state 4-6 were collected for MS2 scans in cross-link discovery mode (both were performed for each sample); monoisotopic precursor selection was enabled and a dynamic exclusion window was set to 30.0 s. Raw files obtained in enumerative mode were analyzed by pFind3 software (Chi et al., 2018) in open search mode and protein modifications inferred by pFind3 and comprising >0.5 % of total protein were included as the variable modifications in pLink2 (Chang et al., 2015) search parameters. pLink2 results were filtered for FDR (<5%), e-value (<10-3), score (<10-2), and abundance (PSMs≥5). Crosslinks were visualized using xiNET (Combe et al., 2015).

### Mass photometry

All data were collected using a Refeyn OneMP mass photometer (Refeyn Ltd) calibrated with bovine serum albumin (66 kDa), beta amylase (224 kDa), and thyroglobulin (660 kDa). Movies were acquired for 6,000 frames (60 s) using AcquireMP software (version 2.4.0) and default settings. Final protein concentrations were empirically determined to achieve ~50 binding events per second. Raw data were converted to frequency distributions using Prism 9 (Graphpad) and a bin size of 15 Da (for shelterin and TRF2/His6MBP-Rap1) (for all other complexes).

For the TRF1 complex, 1 nmol of (TRF1)_2_, 2 nmol POT1/TPP1/GFP-TIN2, and 0 or 1.5 nmol of TeloDNA1 were mixed together and diluted to 250 µL in 20 mM HEPES-KOH pH 7.5, 350 mM NaCl, 0.1 mM TCEP-HCl. After a 2-hr incubation at 4°C, 250 µL of the reconstitution was loaded onto each of two 10-30% glycerol gradients in 20 mM HEPES-KOH pH 7.5, 350 mM NaCl, 0.1 mM TCEP-HCl and centrifuged, fractionated, and analyzed as before. Fractions containing the complex were pooled, concentrated in a Microcon YM-30, buffer exchanged 1:10 into 20 mM HEPES-KOH pH 7.5, 350 mM NaCl, 0.1 mM TCEP-HCl, flash frozen in liquid nitrogen and stored at −80°C. For shelterin, 400 pmol of the TRF1 complex (by absorbance at 488 nm) was mixed with 300 pmol of (TRF2/His_6_MBP-Rap1)_2_ and adjusted to 200 µL with 20 mM HEPES-KOH pH 7.5, 350 mM NaCl, 0.1 mM TCEP-HCl. This sample was dialyzed against 20 mM HEPES-KOH pH 7.5, 150 mM KCl, 0.1 mM TCEP-HCl, 5% v/v glycerol overnight using a 3500 Da cutoff 100-500 µL capacity slide-a-lyzer MINI device (Thermo Fisher) followed by 4 additional hours of dialysis against same buffer lacking glycerol the next morning. The sample was then loaded onto each of two 10-30% glycerol gradients in 20 mM HEPES-KOH pH 7.5, 150 mM KCl, 0.1 mM TCEP-HCl and centrifuged, fractionated, and analyzed by SDS PAGE. Fractions containing the complex were pooled, incubated with 30 µL amylose beads with end-over-end rotation for 3 hrs at 8°C. The beads were then washed three times with 20 mM HEPES-KOH pH 7.5, 150 mM KCl, 0.1 mM TCEP-HCl, and finally eluted with 50 mM maltose in the the same buffer. The eluate was isolated from beads by centrifugation through a 0.22 µm filter using a Spin-X centrifugal filtration device (Corning) and immediately used at a 1:8 final dilution in mass photometry measurements. For the DNA-containing sample, the eluate was first mixed with an equal volume of 60 nM TeloDNA2 (final concentration of 7.5 nM) and incubated on ice for 30 min prior to measurement.

Prior to analysis, samples were thawed, centrifuged at 20,000 x g, and the concentration of the supernatant was measured prior to analysis. Samples were then diluted to 1 µM in 20 mM HEPES-KOH pH 7.5, 300 mM NaCl, 0.1 mM TCEP-HCl, and mixed together if indicated. Immediately prior to measurement, samples were diluted to 4x final concentration in MP buffer (20 mM HEPES-KOH pH 7.5, 150 mM KCl, 0.1 mM TCEP-HCl, filtered three times through 0.22 µm), then 2.5 µL was added to 7.5 µL MP buffer in a sealed well on a glass slide and light scattering was recorded for 1 min.

### Native mass spectrometry

Prior to analysis, aliquots of purified POT1/TPP1/GFP-TIN2 and TRF1 were thawed and treated overnight at 4°C with 50 µg/mL purified lambda protein phosphatase (made in-house) and 1 mM MnCl_2_. For the TRF1 complex, the sample was prepared identically as for mass photometry except using phosphatase treated components. The sample was immediately processed for native mass spectrometry after glycerol exchange.

The samples were buffer-exchanged into 300 mM ammonium acetate (POT1/TPP1/GFP-TIN2) or 400 mM ammonium acetate (TRF1-only and TRF1 complex) solutions at pH 7.5 and with 0.01% Tween-20 using Zeba desalting microspin columns with 40 kDa molecular weight cutoff (Thermo Fisher). Sample concentrations were at least 5 µM for analysis. A 3-µL aliquot of the buffer-exchanged sample was loaded into a gold-coated quartz capillary tip that was prepared in-house. The sample was then electrosprayed into an Exactive Plus EMR instrument (Thermo Fisher) using a modified static nanospray source (Olinares & Chait, 2020; Olinares et al., 2021). The MS parameters used included: spray voltage, 1.2 kV; capillary temperature, 150 °C; S-lens RF level, 200; resolving power, 8,750 or 17,500 at m/z of 200; AGC target, 1 × 10^6^; number of microscans, 5; maximum injection time, 200 ms; in-source dissociation (ISD), 0 V; injection flatapole, 8 V; interflatapole, 4 V (TRF1 trimer complex) or 6 V (for other samples); bent flatapole, 4 V (TRF1 complex) or 6 V (other samples); high energy collision dissociation (HCD), 150-200 V; ultrahigh vacuum pressure, 5.2 − 6.0 × 10^−10^ mbar; total number of scans, at least 100. The instrument mass calibration in positive EMR mode was performed using cesium iodide. The acquired MS spectra were visualized using Thermo Xcalibur Qual Browser (version 4.2.47). Data processing and spectra deconvolution were performed using UniDec version 4.2.0 (Reid et al., 2019; Marty et al., 2015). The general UniDec parameters used included: sample mass every 1.0 Da; smooth charge state distribution, on and peak shape function, Gaussian. No background subtraction was applied for the TRF1-only spectrum and curved background subtraction was set to 10 for spectra from the rest of the samples.

### Antibodies and Western blotting

Polyclonal antibodies were previously generated by inoculating rabbits with the following purified proteins as antigens (produced in insect cells unless otherwise specified): full-length TRF1, full-length TRF2, full-length TIN2, full-length Rap1, and GST-TPP1(87-337) produced in *E. coli*.

Affinity purification of the resulting sera generated primary antibody stocks that were used at 1:2000 dilutions in PBS + 0.05% Tween and 1% v/v milk. A rabbit polyclonal antibody against POT1 was purchased from Proteintech and used at a 1:1000 dilution.

Protein samples were separated on SDS-PAGE and transferred to nitrocellulose membranes. Membranes were blocked for 30 min at room temperature with PBS + 0.05% Tween containing 5% milk prior to incubation with the primary antibody for 60 min at room temperature. Membranes were then washed three times with PBS + 0.05% Tween, incubated with a 1:10,000 dilution of anti-rabbit IgG-HRP (Millipore-Sigma) for 45 min at room temperature in PBS + 0.05% Tween and 1% milk, washed three times, blotted dry, and finally treated with West Pico PLUS luminescence reagent (Thermo Fisher). Membranes were exposed to film for 0.5-5 min, then the film was developed and scanned using the transparency setting of a conventional scanner.

## Author Contributions

John C Zinder, Conceptualization, Formal analysis, Investigation, Methodology, Writing – original draft, Writing – review and editing, Visualization, Funding acquisition; P Dominic B Olinares, Formal analysis, Investigation, Methodology, Writing – review and editing, Visualization; Vladimir Svetlov, Formal analysis, Investigation, Methodology, Writing – review and editing; Martin W Bush, Conceptualization, Writing – review and editing; Evgeny Nudler, Supervision, Writing – review and editing, Funding acquisition; Brian T Chait, Supervision, Writing – review and editing, Funding acquisition; Thomas Walz, Conceptualization, Supervision, Writing – review and editing, Visualization, Funding acquisition; Titia de Lange, Conceptualization, Supervision, Writing – review and editing, Visualization, Funding acquisition.

## Acknowledgements

We would like to thank members of the Walz and de Lange labs for their critical reading of the manuscript and helpful discussions throughout data collection and analysis. We would also like to thank Yixiao Zhang for his assistance in implementing ISAC with GPU acceleration and Lauren Vostal and Tarun Kapoor for training and use of the mass photometry instrument.

## Funding

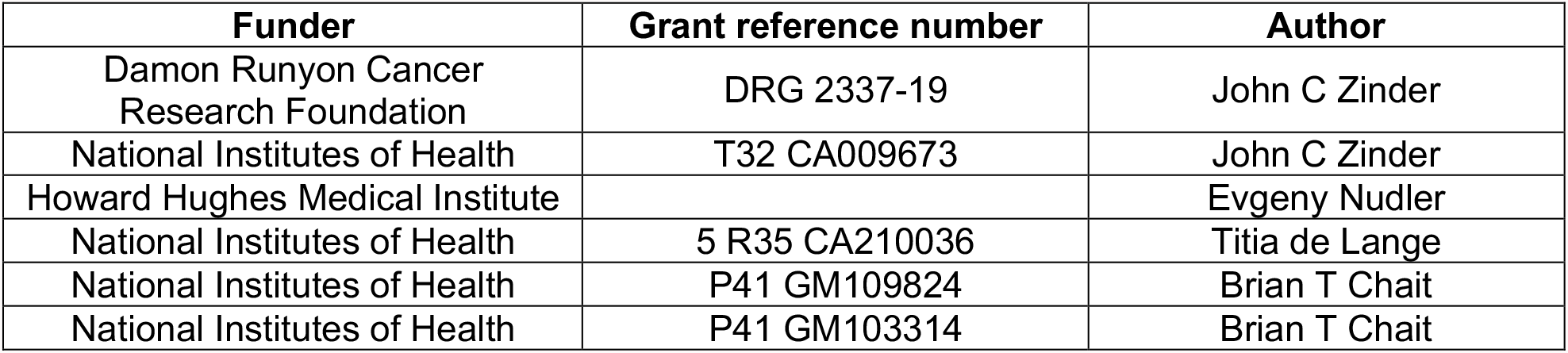

## References

Artandi, S. E., & DePinho, R. A. (2010). Telomeres and telomerase in cancer. Carcinogenesis, 31(1), 9–18. https://doi.org/10.1093/carcin/bgp268

Azzalin, C. M., & Lingner, J. (2015). Telomere functions grounding on TERRA firma. Trends Cell Biol, 25(1), 29–36. https://doi.org/10.1016/j.tcb.2014.08.007

Azzalin, C. M., Reichenbach, P., Khoriauli, L., Giulotto, E., & Lingner, J. (2007). Telomeric repeat containing RNA and RNA surveillance factors at mammalian chromosome ends. Science, 318(5851), 798–801. https://doi.org/10.1126/science.1147182

Baumann, P., & Cech, T. R. (2001). Pot1, the putative telomere end-binding protein in fission yeast and humans. Science, 292(5519), 1171–1175. https://doi.org/10.1126/science.1060036

Bianchi, A., Smith, S., Chong, L., Elias, P., & de Lange, T. (1997). TRF1 is a dimer and bends telomeric DNA. EMBO J, 16(7), 1785–1794. https://doi.org/10.1093/emboj/16.7.1785

Bianchi, A., Stansel, R. M., Fairall, L., Griffith, J. D., Rhodes, D., & de Lange, T. (1999). TRF1 binds a bipartite telomeric site with extreme spatial flexibility. EMBO J, 18(20), 5735–5744. https://doi.org/10.1093/emboj/18.20.5735

Bilaud, T., Brun, C., Ancelin, K., Koering, C. E., Laroche, T., & Gilson, E. (1997). Telomeric localization of TRF2, a novel human telobox protein. Nat Genet, 17(2), 236–239. https://doi.org/10.1038/ng1097-236

Broccoli, D., Smogorzewska, A., Chong, L., & de Lange, T. (1997). Human telomeres contain two distinct Myb-related proteins, TRF1 and TRF2. Nat Genet, 17(2), 231–235. https://doi.org/10.1038/ng1097-231

Chang, C. C., Chow, C. C., Tellier, L. C., Vattikuti, S., Purcell, S. M., & Lee, J. J. (2015). Second-generation PLINK: rising to the challenge of larger and richer datasets. Gigascience, 4, 7. https://doi.org/10.1186/s13742-015-0047-8

Chen, C., Gu, P., Wu, J., Chen, X., Niu, S., Sun, H., Wu, L., Li, N., Peng, J., Shi, S., Fan, C., Huang, M., Wong, C. C., Gong, Q., Kumar-Sinha, C., Zhang, R., Pusztai, L., Rai, R., Chang, S., … Lei, M. (2017). Structural insights into POT1-TPP1 interaction and POT1 C-terminal mutations in human cancer. Nat Commun, 8, 14929. https://doi.org/10.1038/ncomms14929

Chen, L. Y., Redon, S., & Lingner, J. (2012). The human CST complex is a terminator of telomerase activity. Nature, 488(7412), 540–544. https://doi.org/10.1038/nature11269

Chen, Y., Rai, R., Zhou, Z. R., Kanoh, J., Ribeyre, C., Yang, Y., Zheng, H., Damay, P., Wang, F., Tsujii, H., Hiraoka, Y., Shore, D., Hu, H. Y., Chang, S., & Lei, M. (2011). A conserved motif within RAP1 has diversified roles in telomere protection and regulation in different organisms. Nat Struct Mol Biol, 18(2), 213–221. https://doi.org/10.1038/nsmb.1974

Chen, Y., Yang, Y., van Overbeek, M., Donigian, J. R., Baciu, P., de Lange, T., & Lei, M. (2008). A shared docking motif in TRF1 and TRF2 used for differential recruitment of telomeric proteins. Science, 319(5866), 1092–1096. https://doi.org/10.1126/science.1151804

Chi, H., Liu, C., Yang, H., Zeng, W. F., Wu, L., Zhou, W. J., Wang, R. M., Niu, X. N., Ding, Y. H., Zhang, Y., Wang, Z. W., Chen, Z. L., Sun, R. X., Liu, T., Tan, G. M., Dong, M. Q., Xu, P., Zhang, P. H., & He, S. M. (2018). Comprehensive identification of peptides in tandem mass spectra using an efficient open search engine. Nat Biotechnol. https://doi.org/10.1038/nbt.4236

Chong, L., van Steensel, B., Broccoli, D., Erdjument-Bromage, H., Hanish, J., Tempst, P., & de Lange, T. (1995). A human telomeric protein. Science, 270(5242), 1663–1667. https://doi.org/10.1126/science.270.5242.1663

Combe, C. W., Fischer, L., & Rappsilber, J. (2015). xiNET: cross-link network maps with residue resolution. Mol Cell Proteomics, 14(4), 1137–1147. https://doi.org/10.1074/mcp.O114.042259

Court, R., Chapman, L., Fairall, L., & Rhodes, D. (2005). How the human telomeric proteins TRF1 and TRF2 recognize telomeric DNA: a view from high-resolution crystal structures. EMBO Rep, 6(1), 39–45. https://doi.org/10.1038/sj.embor.7400314

de Lange, T. (2005). Shelterin: the protein complex that shapes and safeguards human telomeres. Genes Dev, 19(18), 2100–2110. https://doi.org/10.1101/gad.1346005

de Lange, T. (2018). Shelterin-Mediated Telomere Protection. Annu Rev Genet, 52, 223–247. https://doi.org/10.1146/annurev-genet-032918-021921

Denchi, E. L., & de Lange, T. (2007). Protection of telomeres through independent control of ATM and ATR by TRF2 and POT1. Nature, 448(7157), 1068–1071. https://doi.org/10.1038/nature06065

Doksani, Y., Wu, J. Y., de Lange, T., & Zhuang, X. (2013). Super-resolution fluorescence imaging of telomeres reveals TRF2-dependent T-loop formation. Cell, 155(2), 345–356. https://doi.org/10.1016/j.cell.2013.09.048

Erdel, F., Kratz, K., Willcox, S., Griffith, J. D., Greene, E. C., & de Lange, T. (2017). Telomere Recognition and Assembly Mechanism of Mammalian Shelterin. Cell Rep, 18(1), 41–53. https://doi.org/10.1016/j.celrep.2016.12.005

Fairall, L., Chapman, L., Moss, H., de Lange, T., & Rhodes, D. (2001). Structure of the TRFH dimerization domain of the human telomeric proteins TRF1 and TRF2. Mol Cell, 8(2), 351–361. https://doi.org/10.1016/s1097-2765(01)00321-5

Frescas, D., & de Lange, T. (2014). TRF2-tethered TIN2 can mediate telomere protection by TPP1/POT1. Mol Cell Biol, 34(7), 1349–1362. https://doi.org/10.1128/MCB.01052-13

Ghanim, G. E., Fountain, A. J., van Roon, A. M., Rangan, R., Das, R., Collins, K., & Nguyen, T. H. D. (2021). Structure of human telomerase holoenzyme with bound telomeric DNA. Nature, 593(7859), 449–453. https://doi.org/10.1038/s41586-021-03415-4

Gibson, D. G., Young, L., Chuang, R. Y., Venter, J. C., Hutchison, C. A., & Smith, H. O. (2009). Enzymatic assembly of DNA molecules up to several hundred kilobases. Nat Methods, 6(5), 343–345. https://doi.org/10.1038/nmeth.1318

Gong, Y., & de Lange, T. (2010). A Shld1-controlled POT1a provides support for repression of ATR signaling at telomeres through RPA exclusion. Mol Cell, 40(3), 377–387. https://doi.org/10.1016/j.molcel.2010.10.016

Greider, C. W., & Blackburn, E. H. (1985). Identification of a specific telomere terminal transferase activity in Tetrahymena extracts. Cell, 43(2 Pt 1), 405–413. https://doi.org/10.1016/0092-8674(85)90170-9

Griffith, J. D., Comeau, L., Rosenfield, S., Stansel, R. M., Bianchi, A., Moss, H., & de Lange, T. (1999). Mammalian telomeres end in a large duplex loop. Cell, 97(4), 503–514. https://doi.org/10.1016/s0092-8674(00)80760-6

Hanaoka, S., Nagadoi, A., Yoshimura, S., Aimoto, S., Li, B., de Lange, T., & Nishimura, Y. (2001). NMR structure of the hRap1 Myb motif reveals a canonical three-helix bundle lacking the positive surface charge typical of Myb DNA-binding domains. J Mol Biol, 312(1), 167–175. https://doi.org/10.1006/jmbi.2001.4924

Hockemeyer, D., Daniels, J. P., Takai, H., & de Lange, T. (2006). Recent expansion of the telomeric complex in rodents: Two distinct POT1 proteins protect mouse telomeres. Cell, 126(1), 63–77. https://doi.org/10.1016/j.cell.2006.04.044

Houghtaling, B. R., Cuttonaro, L., Chang, W., & Smith, S. (2004). A dynamic molecular link between the telomere length regulator TRF1 and the chromosome end protector TRF2. Curr Biol, 14(18), 1621–1631. https://doi.org/10.1016/j.cub.2004.08.052

Hu, C., Rai, R., Huang, C., Broton, C., Long, J., Xu, Y., Xue, J., Lei, M., Chang, S., & Chen, Y. (2017). Structural and functional analyses of the mammalian TIN2-TPP1-TRF2 telomeric complex. Cell Res, 27(12), 1485–1502. https://doi.org/10.1038/cr.2017.144

Kim, S. H., Kaminker, P., & Campisi, J. (1999). TIN2, a new regulator of telomere length in human cells. Nat Genet, 23(4), 405–412. https://doi.org/10.1038/70508

König, P., Fairall, L., & Rhodes, D. (1998). Sequence-specific DNA recognition by the myb-like domain of the human telomere binding protein TRF1: a model for the protein-DNA complex. Nucleic Acids Res, 26(7), 1731–1740. https://doi.org/10.1093/nar/26.7.1731

Kratz, K., & de Lange, T. (2018). Protection of telomeres 1 proteins POT1a and POT1b can repress ATR signaling by RPA exclusion, but binding to CST limits ATR repression by POT1b. J Biol Chem, 293(37), 14384–14392. https://doi.org/10.1074/jbc.RA118.004598

Lam, Y. C., Akhter, S., Gu, P., Ye, J., Poulet, A., Giraud-Panis, M. J., Bailey, S. M., Gilson, E., Legerski, R. J., & Chang, S. (2010). SNMIB/Apollo protects leading-strand telomeres against NHEJ-mediated repair. EMBO J, 29(13), 2230–2241. https://doi.org/10.1038/emboj.2010.58

Lei, M., Podell, E. R., & Cech, T. R. (2004). Structure of human POT1 bound to telomeric single-stranded DNA provides a model for chromosome end-protection. Nat Struct Mol Biol, 11(12), 1223–1229. https://doi.org/10.1038/nsmb867

Lenain, C., Bauwens, S., Amiard, S., Brunori, M., Giraud-Panis, M. J., & Gilson, E. (2006). The Apollo 5’ exonuclease functions together with TRF2 to protect telomeres from DNA repair. Curr Biol, 16(13), 1303–1310. https://doi.org/10.1016/j.cub.2006.05.021

Lim, C. J., & Cech, T. R. (2021). Shaping human telomeres: from shelterin and CST complexes to telomeric chromatin organization. Nat Rev Mol Cell Biol, 22(4), 283–298. https://doi.org/10.1038/s41580-021-00328-y

Lim, C. J., Zaug, A. J., Kim, H. J., & Cech, T. R. (2017). Reconstitution of human shelterin complexes reveals unexpected stoichiometry and dual pathways to enhance telomerase processivity. Nat Commun, 8(1), 1075. https://doi.org/10.1038/s41467-017-01313-w

Lingner, J., Hughes, T. R., Shevchenko, A., Mann, M., Lundblad, V., & Cech, T. R. (1997). Reverse transcriptase motifs in the catalytic subunit of telomerase. Science, 276(5312), 561–567. https://doi.org/10.1126/science.276.5312.561

Liu, D., O’Connor, M. S., Qin, J., & Songyang, Z. (2004a). Telosome, a mammalian telomere-associated complex formed by multiple telomeric proteins. J Biol Chem, 279(49), 51338–51342. https://doi.org/10.1074/jbc.M409293200

Liu, D., Safari, A., O’Connor, M. S., Chan, D. W., Laegeler, A., Qin, J., & Songyang, Z. (2004b). PTOP interacts with POT1 and regulates its localization to telomeres. Nat Cell Biol, 6(7), 673–680. https://doi.org/10.1038/ncb1142

Liu, P., Zupa, E., Neuner, A., Böhler, A., Loerke, J., Flemming, D., Ruppert, T., Rudack, T., Peter, C., Spahn, C., Gruss, O. J., Pfeffer, S., & Schiebel, E. (2020). Insights into the assembly and activation of the microtubule nucleator γ-TuRC. Nature, 578(7795), 467–471. https://doi.org/10.1038/s41586-019-1896-6

Maciejowski, J., & de Lange, T. (2017). Telomeres in cancer: tumour suppression and genome instability. Nat Rev Mol Cell Biol, 18(3), 175–186. https://doi.org/10.1038/nrm.2016.171

Martínez, P., Gómez-López, G., García, F., Mercken, E., Mitchell, S., Flores, J. M., de Cabo, R., & Blasco, M. A. (2013). RAP1 protects from obesity through its extratelomeric role regulating gene expression. Cell Rep, 3(6), 2059–2074. https://doi.org/10.1016/j.celrep.2013.05.030

Marty, M. T., Baldwin, A. J., Marklund, E. G., Hochberg, G. K., Benesch, J. L., & Robinson, C. V. (2015). Bayesian deconvolution of mass and ion mobility spectra: from binary interactions to polydisperse ensembles. Anal Chem, 87(8), 4370–4376. https://doi.org/10.1021/acs.analchem.5b00140

Meyer, A., Dierks, K., Hussein, R., Brillet, K., Brognaro, H., & Betzel, C. (2015). Systematic analysis of protein-detergent complexes applying dynamic light scattering to optimize solutions for crystallization trials. Acta Crystallogr F Struct Biol Commun, 71(Pt 1), 75–81. https://doi.org/10.1107/S2053230X14027149

Myler, L. R., Kinzig, C. G., Sasi, N. K., Zakusilo, G., Cai, S. W., & de Lange, T. (2021). The evolution of metazoan shelterin. Genes Dev, 35(23-24), 1625–1641. https://doi.org/10.1101/gad.348835.121

Nandakumar, J., Bell, C. F., Weidenfeld, I., Zaug, A. J., Leinwand, L. A., & Cech, T. R. (2012). The TEL patch of telomere protein TPP1 mediates telomerase recruitment and processivity. Nature, 492(7428), 285–289. https://doi.org/10.1038/nature11648

Nelson, N. D., & Bertuch, A. A. (2012). Dyskeratosis congenita as a disorder of telomere maintenance. Mutat Res, 730(1-2), 43–51. https://doi.org/10.1016/j.mrfmmm.2011.06.008

Olinares, P. D. B., & Chait, B. T. (2020). Native Mass Spectrometry Analysis of Affinity-Captured Endogenous Yeast RNA Exosome Complexes. Methods Mol Biol, 2062, 357–382. https://doi.org/10.1007/978-1-4939-9822-7_17

Olinares, P. D. B., Kang, J. Y., Llewellyn, E., Chiu, C., Chen, J., Malone, B., Saecker, R. M., Campbell, E. A., Darst, S. A., & Chait, B. T. (2021). Native Mass Spectrometry-Based Screening for Optimal Sample Preparation in Single-Particle Cryo-EM. Structure, 29(2), 186–195.e6. https://doi.org/10.1016/j.str.2020.11.001

Reid, D. J., Diesing, J. M., Miller, M. A., Perry, S. M., Wales, J. A., Montfort, W. R., & Marty, M. T. (2019). MetaUniDec: High-Throughput Deconvolution of Native Mass Spectra. J Am Soc Mass Spectrom, 30(1), 118–127. https://doi.org/10.1007/s13361-018-1951-9

Rice, C., Shastrula, P. K., Kossenkov, A. V., Hills, R., Baird, D. M., Showe, L. C., Doukov, T., Janicki, S., & Skordalakes, E. (2017). Structural and functional analysis of the human POT1-TPP1 telomeric complex. Nat Commun, 8, 14928. https://doi.org/10.1038/ncomms14928

Sarek, G., Vannier, J. B., Panier, S., Petrini, J. H. J., & Boulton, S. J. (2015). TRF2 recruits RTEL1 to telomeres in S phase to promote t-loop unwinding. Mol Cell, 57(4), 622–635. https://doi.org/10.1016/j.molcel.2014.12.024

Savage, S. A., & Bertuch, A. A. (2010). The genetics and clinical manifestations of telomere biology disorders. Genet Med, 12, 753–764. https://doi.org/10.1097/GIM.0b013e3181f415b5

Savage, S. A., Giri, N., Jessop, L., Pike, K., Plona, T., Burdett, L., & Alter, B. P. (2011). Sequence analysis of the shelterin telomere protection complex genes in dyskeratosis congenita. J Med Genet, 48(4), 285–288. https://doi.org/10.1136/jmg.2010.082727

Sfeir, A., Kabir, S., van Overbeek, M., Celli, G. B., & de Lange, T. (2010). Loss of Rap1 induces telomere recombination in the absence of NHEJ or a DNA damage signal. Science, 327(5973), 1657–1661. https://doi.org/10.1126/science.1185100

Sfeir, A., Kosiyatrakul, S. T., Hockemeyer, D., MacRae, S. L., Karlseder, J., Schildkraut, C. L., & de Lange, T. (2009). Mammalian telomeres resemble fragile sites and require TRF1 for efficient replication. Cell, 138(1), 90–103. https://doi.org/10.1016/j.cell.2009.06.021

Sonn-Segev, A., Belacic, K., Bodrug, T., Young, G., VanderLinden, R. T., Schulman, B. A., Schimpf, J., Friedrich, T., Dip, P. V., Schwartz, T. U., Bauer, B., Peters, J. M., Struwe, W. B., Benesch, J. L. P., Brown, N. G., Haselbach, D., & Kukura, P. (2020). Quantifying the heterogeneity of macromolecular machines by mass photometry. Nat Commun, 11(1), 1772. https://doi.org/10.1038/s41467-020-15642-w

Takai, K. K., Hooper, S., Blackwood, S., Gandhi, R., & de Lange, T. (2010). In vivo stoichiometry of shelterin components. J Biol Chem, 285(2), 1457–1467. https://doi.org/10.1074/jbc.M109.038026

Takai, K. K., Kibe, T., Donigian, J. R., Frescas, D., & de Lange, T. (2011). Telomere protection by TPP1/POT1 requires tethering to TIN2. Mol Cell, 44(4), 647–659. https://doi.org/10.1016/j.molcel.2011.08.043

Tang, G., Peng, L., Baldwin, P. R., Mann, D. S., Jiang, W., Rees, I., & Ludtke, S. J. (2007). EMAN2: an extensible image processing suite for electron microscopy. J Struct Biol, 157(1), 38–46. https://doi.org/10.1016/j.jsb.2006.05.009

Telomeres, M. R. C., Haycock, P. C., Burgess, S., Nounu, A., Zheng, J., Okoli, G. N., Bowden, J., Wade, K. H., Timpson, N. J., Evans, D. M., Willeit, P., Aviv, A., Gaunt, T. R., Hemani, G., Mangino, M., Ellis, H. P., Kurian, K. M., Pooley, K. A., Eeles, R. A., … Davey Smith, G. (2017). Association Between Telomere Length and Risk of Cancer and Non-Neoplastic Diseases: A Mendelian Randomization Study. JAMA Oncol, 3(5), 636–651. https://doi.org/10.1001/jamaoncol.2016.5945

Timashev, L. A., & de Lange, T. (2020). Characterization of t-loop formation by TRF2. Nucleus, 11(1), 164–177. https://doi.org/10.1080/19491034.2020.1783782

van Overbeek, M., & de Lange, T. (2006). Apollo, an Artemis-related nuclease, interacts with TRF2 and protects human telomeres in S phase. Curr Biol, 16(13), 1295–1302. https://doi.org/10.1016/j.cub.2006.05.022

Wan, F., Ding, Y., Zhang, Y., Wu, Z., Li, S., Yang, L., Yan, X., Lan, P., Li, G., Wu, J., & Lei, M. (2021). Zipper head mechanism of telomere synthesis by human telomerase. Cell Res, 31(12), 1275–1290. https://doi.org/10.1038/s41422-021-00586-7

Wan, M., Qin, J., Songyang, Z., & Liu, D. (2009). OB fold-containing protein 1 (OBFC1), a human homolog of yeast Stn1, associates with TPP1 and is implicated in telomere length regulation. J Biol Chem, 284(39), 26725–26731. https://doi.org/10.1074/jbc.M109.021105

Wang, F., Podell, E. R., Zaug, A. J., Yang, Y., Baciu, P., Cech, T. R., & Lei, M. (2007). The POT1-TPP1 telomere complex is a telomerase processivity factor. Nature, 445(7127), 506–510. https://doi.org/10.1038/nature05454

Weissmann, F., Petzold, G., VanderLinden, R., Huis In ‘t Veld, P. J., Brown, N. G., Lampert, F., Westermann, S., Stark, H., Schulman, B. A., & Peters, J. M. (2016). biGBac enables rapid gene assembly for the expression of large multisubunit protein complexes. Proc Natl Acad Sci U S A, 113(19), E2564–9. https://doi.org/10.1073/pnas.1604935113

Wieczorek, M., Urnavicius, L., Ti, S. C., Molloy, K. R., Chait, B. T., & Kapoor, T. M. (2020). Asymmetric Molecular Architecture of the Human γ-Tubulin Ring Complex. Cell, 180(1), 165–175.e16. https://doi.org/10.1016/j.cell.2019.12.007

Wu, L., Multani, A. S., He, H., Cosme-Blanco, W., Deng, Y., Deng, J. M., Bachilo, O., Pathak, S., Tahara, H., Bailey, S. M., Deng, Y., Behringer, R. R., & Chang, S. (2006). Pot1 deficiency initiates DNA damage checkpoint activation and aberrant homologous recombination at telomeres. Cell, 126(1), 49–62. https://doi.org/10.1016/j.cell.2006.05.037

Wu, P., Takai, H., & de Lange, T. (2012). Telomeric 3’ overhangs derive from resection by Exo1 and Apollo and fill-in by POT1b-associated CST. Cell, 150(1), 39–52. https://doi.org/10.1016/j.cell.2012.05.026

Wu, P., van Overbeek, M., Rooney, S., & de Lange, T. (2010). Apollo contributes to G overhang maintenance and protects leading-end telomeres. Mol Cell, 39(4), 606–617. https://doi.org/10.1016/j.molcel.2010.06.031

Yang, Z., Fang, J., Chittuluru, J., Asturias, F. J., & Penczek, P. A. (2012). Iterative stable alignment and clustering of 2D transmission electron microscope images. Structure, 20(2), 237–247. https://doi.org/10.1016/j.str.2011.12.007

Yang, Z., Takai, K. K., Lovejoy, C. A., & de Lange, T. (2020). Break-induced replication promotes fragile telomere formation. Genes Dev, 34(19-20), 1392–1405. https://doi.org/10.1101/gad.328575.119

Ye, J. Z., Donigian, J. R., van Overbeek, M., Loayza, D., Luo, Y., Krutchinsky, A. N., Chait, B. T., & de Lange, T. (2004a). TIN2 binds TRF1 and TRF2 simultaneously and stabilizes the TRF2 complex on telomeres. J Biol Chem, 279(45), 47264–47271. https://doi.org/10.1074/jbc.M409047200

Ye, J. Z., Hockemeyer, D., Krutchinsky, A. N., Loayza, D., Hooper, S. M., Chait, B. T., & de Lange, T. (2004b). POT1-interacting protein PIP1: a telomere length regulator that recruits POT1 to the TIN2/TRF1 complex. Genes Dev, 18(14), 1649–1654. https://doi.org/10.1101/gad.1215404

Young, G., Hundt, N., Cole, D., Fineberg, A., Andrecka, J., Tyler, A., Olerinyova, A., Ansari, A., Marklund, E. G., Collier, M. P., Chandler, S. A., Tkachenko, O., Allen, J., Crispin, M., Billington, N., Takagi, Y., Sellers, J. R., Eichmann, C., Selenko, P., … Kukura, P. (2018). Quantitative mass imaging of single biological macromolecules. Science, 360(6387), 423–427. https://doi.org/10.1126/science.aar5839

Zhong, F. L., Batista, L. F., Freund, A., Pech, M. F., Venteicher, A. S., & Artandi, S. E. (2012). TPP1 OB-fold domain controls telomere maintenance by recruiting telomerase to chromosome ends. Cell, 150(3), 481–494. https://doi.org/10.1016/j.cell.2012.07.012

Zimmermann, M., Kibe, T., Kabir, S., & de Lange, T. (2014). TRF1 negotiates TTAGGG repeat-associated replication problems by recruiting the BLM helicase and the TPP1/POT1 repressor of ATR signaling. Genes Dev, 28(22), 2477–2491. https://doi.org/10.1101/gad.251611.114

Zivanov, J., Nakane, T., Forsberg, B. O., Kimanius, D., Hagen, W. J., Lindahl, E., & Scheres, S. H. (2018). New tools for automated high-resolution cryo-EM structure determination in RELION-3. Elife, 7, e42166. https://doi.org/10.7554/eLife.42166

