## Supplemental Table for "Shelterin is a Dimeric Complex with Extensive Structural Heterogeneity"

*To whom correspondence should be addressed.

Zinder et al. Figure 1 - figure supplement 1

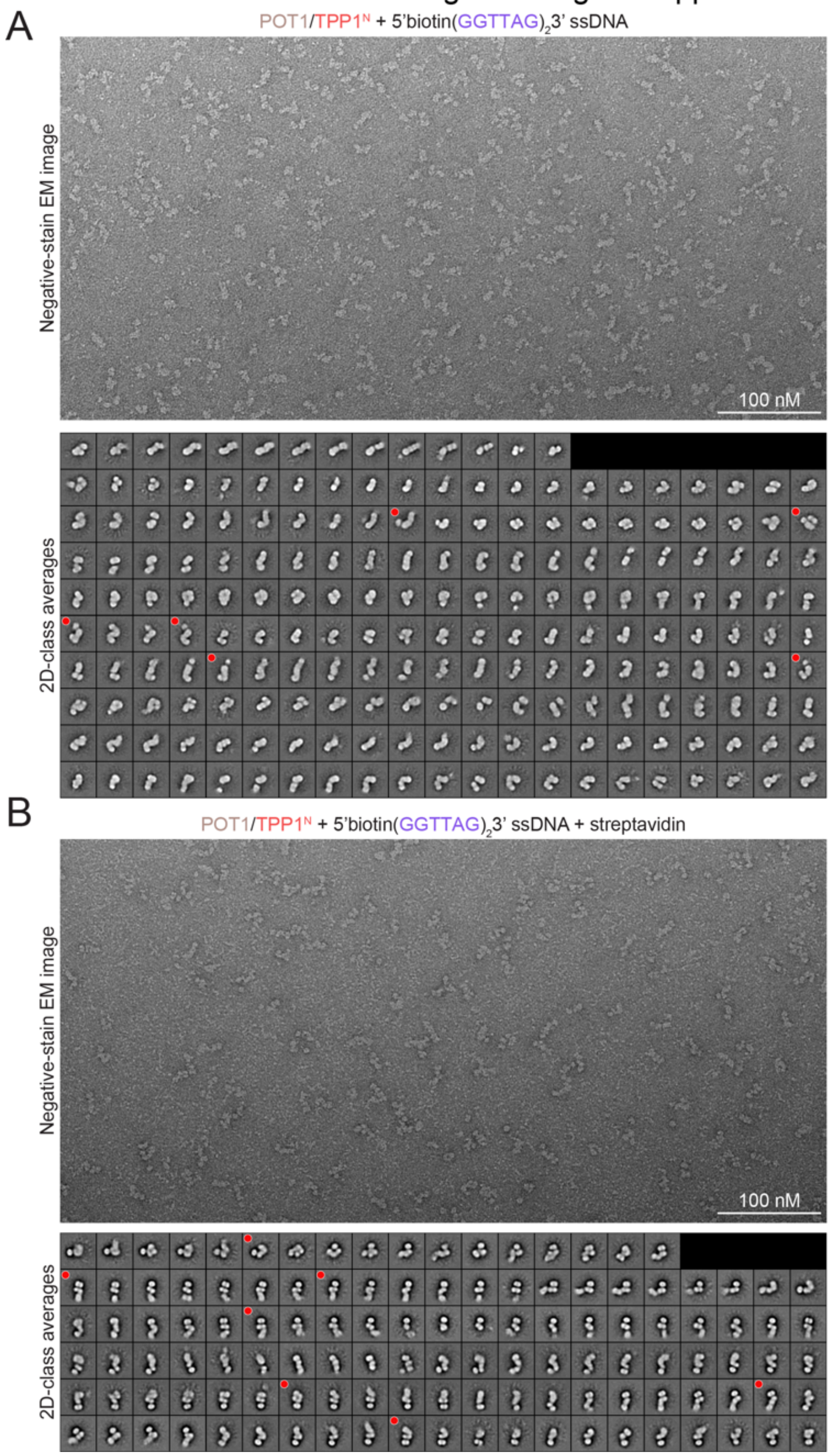


**Figure 1—figure supplement 1. Negative-stain EM of ssDNA bound POT1/TPP1$^N$.**
**(A)** Raw negative-stain EM image and 2D-class averages of POT1/TPP1$^N$ bound to 5'BiotinGGTTAGGGTTAG3' ssDNA. Classes selected for display in Figure 1D are indicated with red dots and may have been rotated and/or reflected. **(B)** Raw negative-stain EM image and 2D-class averages of POT1/TPP1$^N$ bound to streptavidin-5'BiotinGGTTAGGGTTAG3' ssDNA. Classes selected for display in Figure 1E are indicated with red dots.

Zinder et al. Figure 1 - figure supplement 2

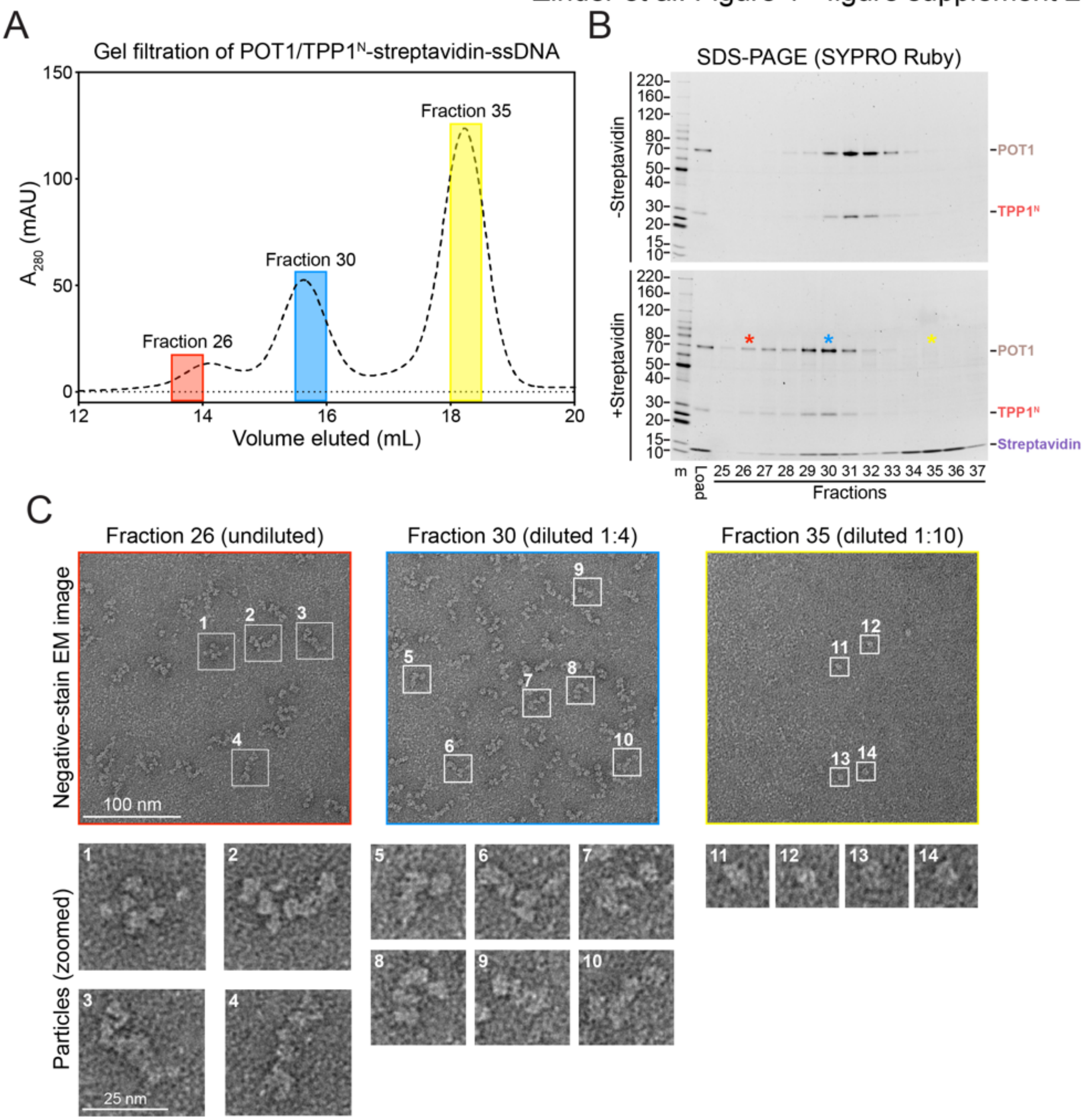


**Figure 1—figure supplement 2**. **Gel filtration and negative-stain EM analysis of POT1/TPP1$^{N}$ bound to streptavidin-ssDNA**

**(A)** Superose 6 increase 10/300 GL gel-filtration trace for absorbance at 280 nm of streptavidin-5'BiotinGGTTAGGGTTAG3' ssDNA from Figure 1D with selected fractions highlighted. **(B)** SDS-PAGE of fractions from gel filtration. Gel is 8-16% polyacrylamide Tris-glycine and stained with SYPRO Ruby. Asterisks correspond to the selected fractions in panel A. Molecular-weight markers are shown in kDa. **(C)** Raw negative-stain EM images of the indicated fractions with selected particles highlighted and zoomed below.

Zinder et al. Figure 2 - figure supplement 1

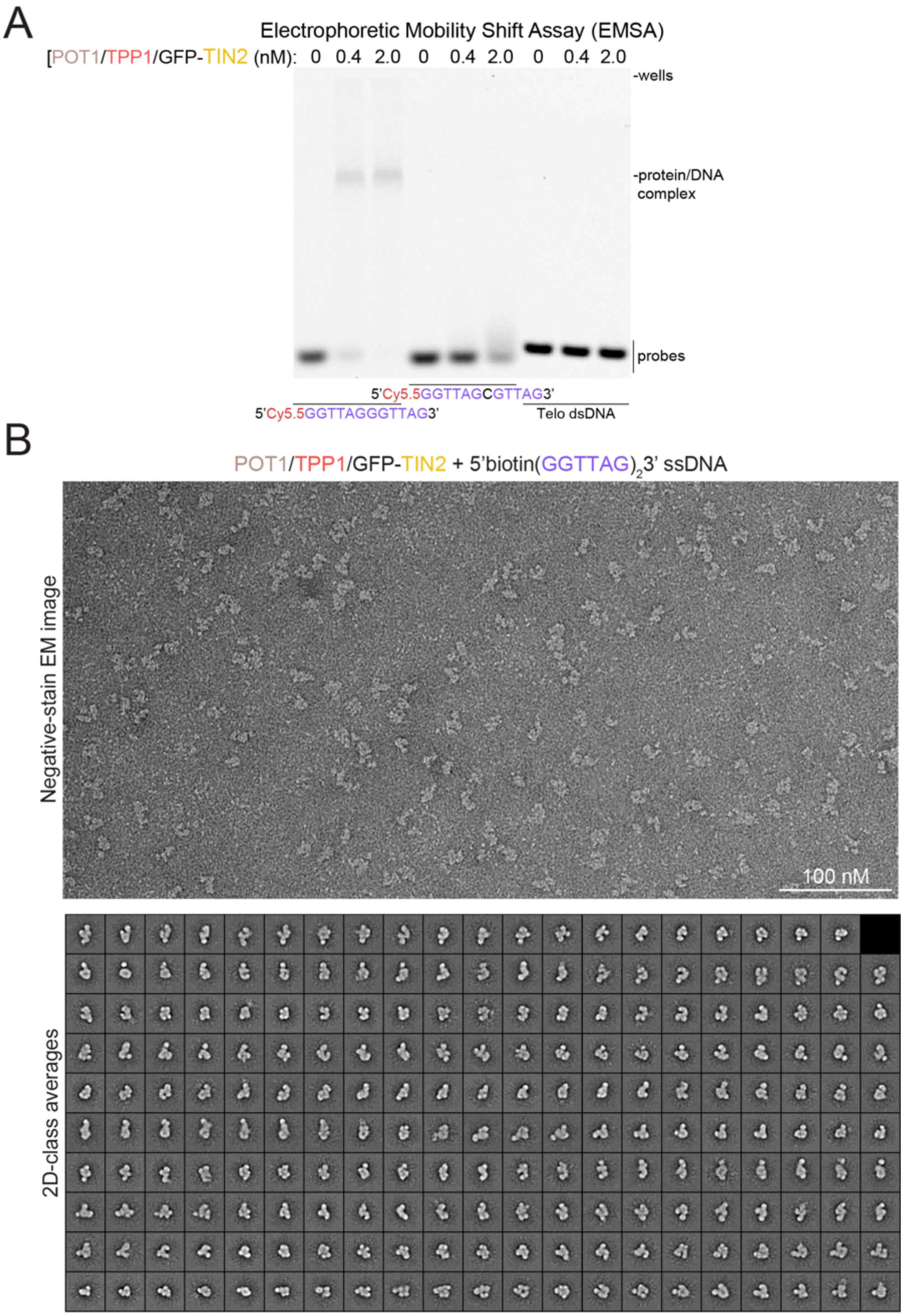


**Figure 2—figure supplement 1. DNA Binding and negative-stain EM analysis of POT1/TPP1/GFP-TIN2**

**(A)** DNA-binding activity of POT1/TPP1/GFP-TIN2 on telomeric and mutant DNAs. Protein concentrations and ssDNA sequences are indicated. 'Telo dsDNA' with sequence 5'CATCAATAGGGTTCATCCTAGGGTTGTACTG3' was labeled with Cy5.5 dUTP by Klenow polymerase. Probe concentration is 0.25 nM. **(B)** Raw negative-stain EM image and 2D-class averages of POT1/TPP1/GFP-TIN2 bound to 5'BiotinGGTTAGGGTTAG3' ssDNA.

Zinder et al. Figure 2 - figure supplement 2

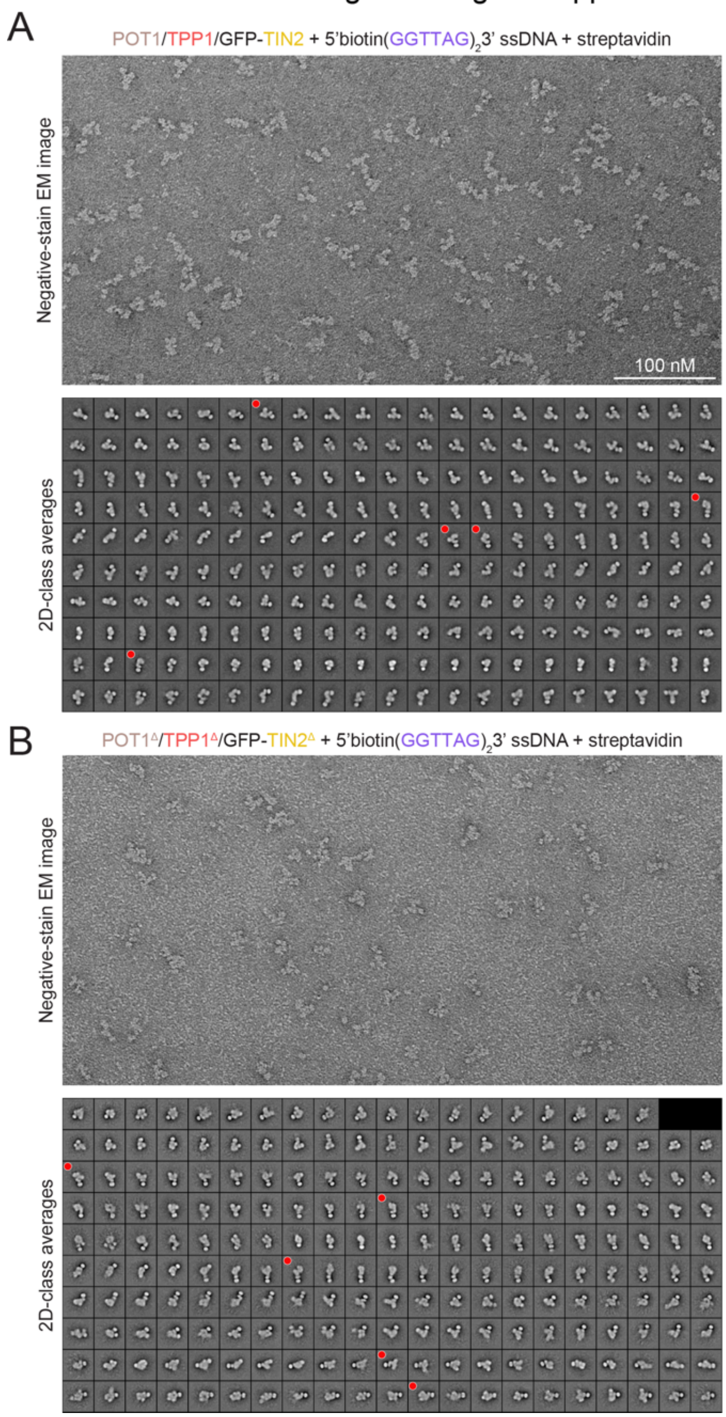


**Figure 2—figure supplement 2. Negative-stain EM of wild-type and 3x∆ POT1/TPP1/GFP-TIN2 bound to streptavidin-ssDNA**

**(A)** Raw negative-stain EM image and 2D-class averages of POT1/TPP1/GFP-TIN2 bound to streptavidin-5'BiotinGGTTAGGGTTAG3' ssDNA. Classes selected for display in Figure 2D are indicated with red dots and may have been rotated and/or reflected. **(B)** Raw negative-stain EM image and 2D-class averages of POT1$^{\Delta}$/TPP1$^{\Delta}$/GFP-TIN2$^{\Delta}$ bound to streptavidin-5'biotinGGTTAGGGTTAG3' ssDNA. Classes selected for display in Figure 2D are indicated with red dots and may have been rotated and/or reflected.

Zinder et al. Figure 3 - figure supplement 1

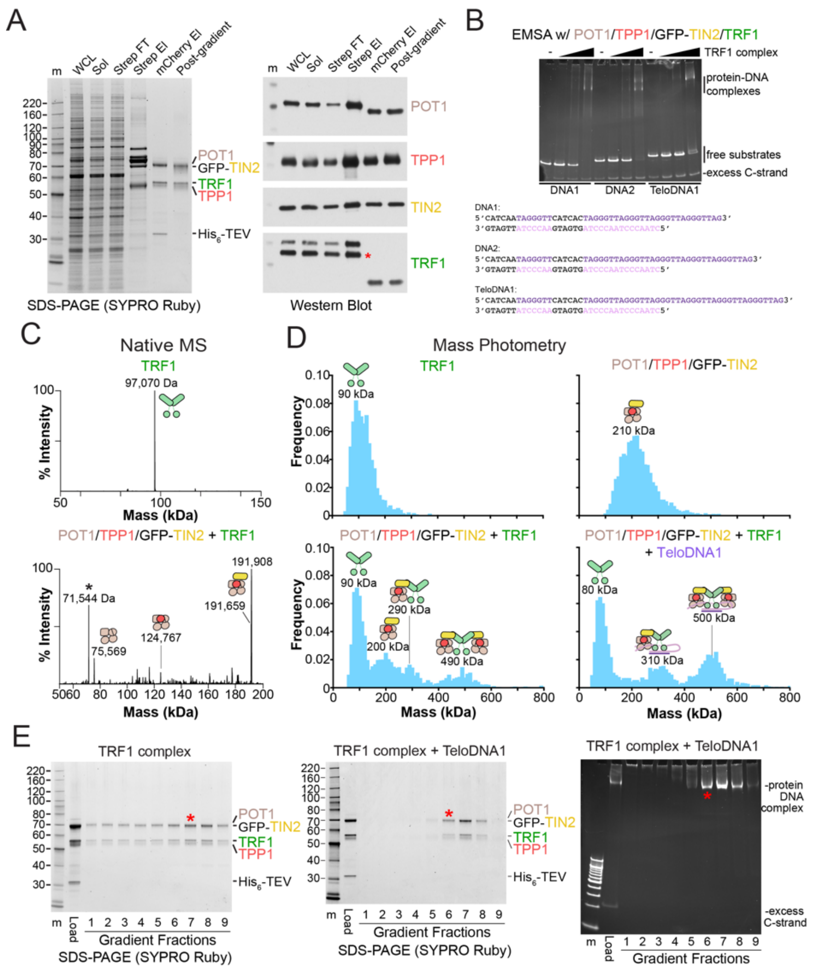


**Figure 3—figure supplement 1. Biochemical analyses and reconstitution of the TRF1 complex**

**(A)** SDS-PAGE and Western blot summary of TRF1 complex purification. Gel is 8-16% polyacrylamide Tris-glycine and stained with SYPRO Ruby. Molecular-weight markers are shown in kDa. WCL: whole-cell lysate; Sol: soluble extract; FT: flow-through; El: eluate. Asterisk corresponds to an unknown species derived from the mCherry tag. **(B)** Native MS of purified and phosphatase-treated TRF1 and POT1/TPP1/GFP-TIN2 samples. The measured masses (in Da) are indicated along with cartoons that correspond to complexes. Asterisk indicates a species that likely corresponds to HSP70 contamination. **(C)** Qualitative EMSA of the TRF1 complex post mCherry elution binding to the indicated DNAs. DNAs were present at 50 nM and the approximate concentration of the TRF1 complex was determined by GFP absorbance at 488 nm. Gel is 4-20% polyacrylamide TBE and stained with SYBR Gold. T1comp: TRF1 complex **(D)** Mass photometry of TRF1 complex components and reconstituted TRF1 complex. Final protein concentrations were 50 nM $(TRF1)_2$, 25 nM POT1/TPP1/GFP-TIN2, 6 nM $(TRF1)_2$ + 12 nM POT1/TPP1/GFP-TIN2, and 6 nM $(TRF1)_2$ + 12 nM POT1/TPP1/GFP-TIN2 + 6 nM TeloDNA1. Peak maxima are indicated along with cartoons that correspond to complexes of that molecular weight to within ~10%. **(E)** SDS-PAGE of fractions from glycerol-gradient purification. Lower number fractions have higher glycerol concentrations. SDS-PAGE gel is 8-16% polyacrylamide Tris-glycine and stained with SYPRO Ruby and native TBE gel is 4-20% polyacrylamide TBE and stained with SYBR Gold. Asterisks indicate fractions that were used for negative-stain EM analysis.

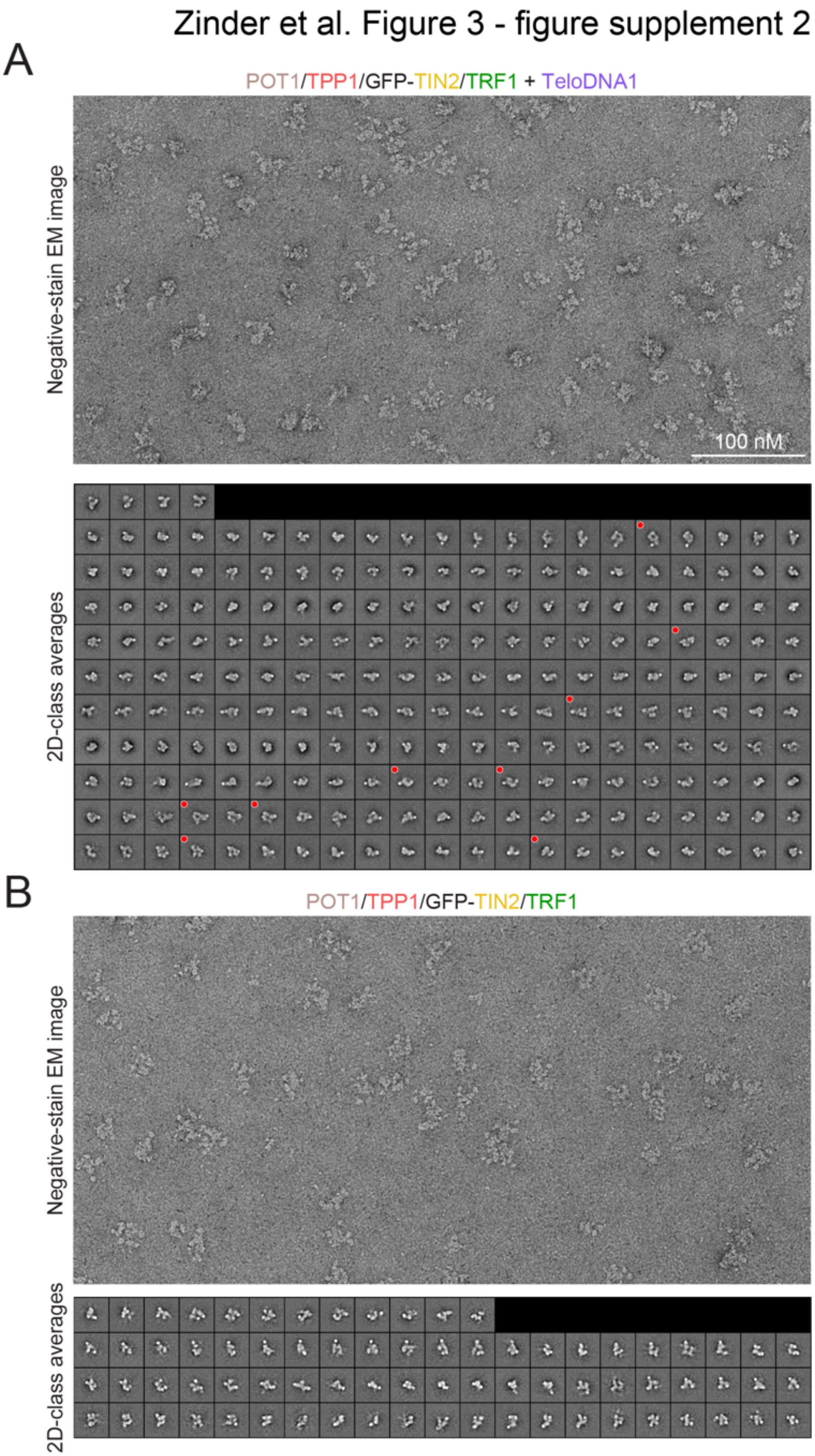


**Figure 3—figure supplement 2. Negative-stain EM of the TRF1 complex**
**(A)** Raw negative-stain EM image and 2D-class averages of the TRF1 complex bound to TeloDNA1 ssDNA. Classes selected for display in Figure 3F are indicated with red dots. **(B)** Raw negative-stain EM image and 2D-class averages of the TRF1 complex.

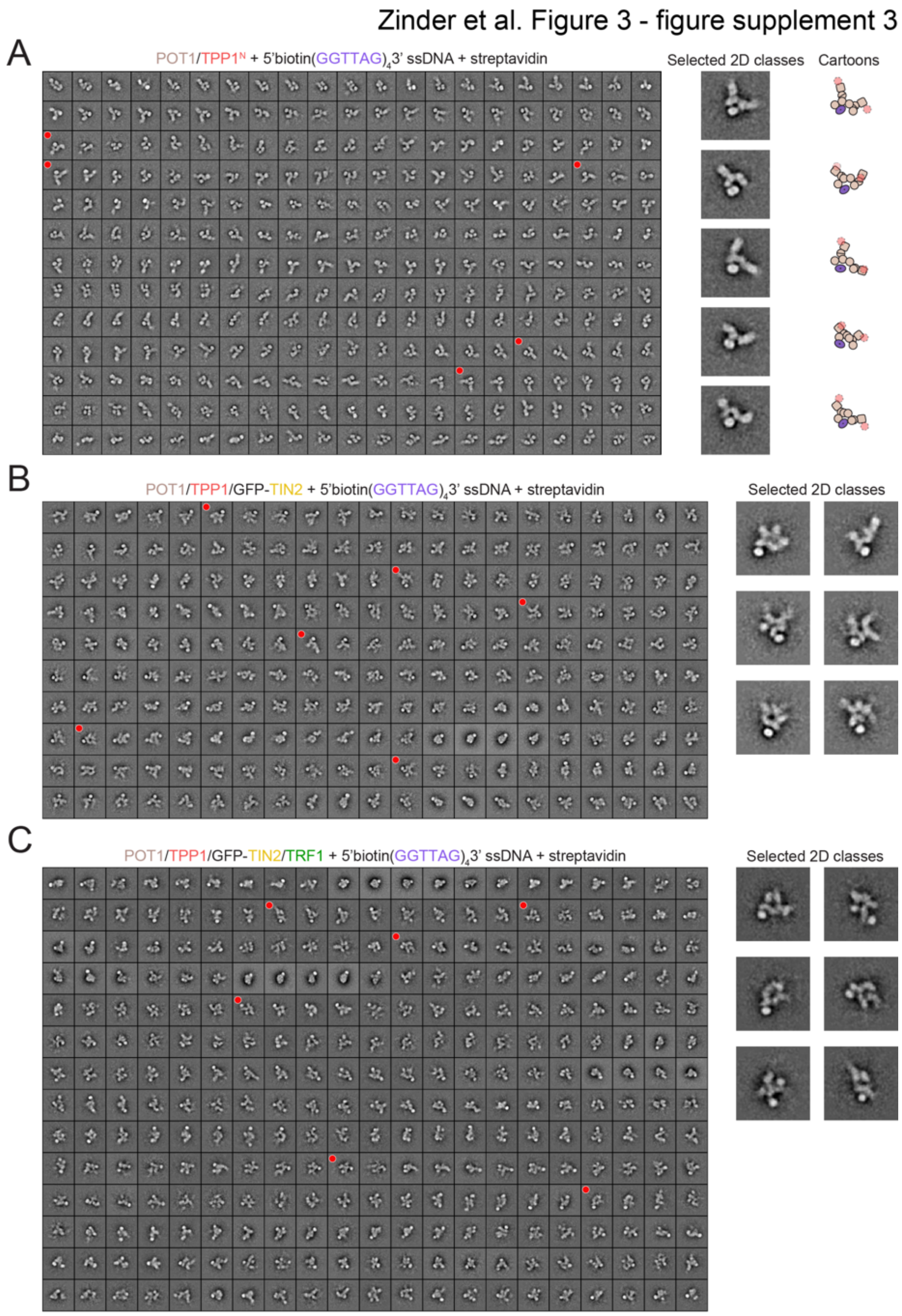


**Figure 3—figure supplement 3. Negative-stain EM of shelterin subcomplexes bound to DNA containing two POT1-binding sites**

**(A)** 2D-class averages of POT1/TPP1$^N$ bound to streptavidin-5'Biotin(GGTTAG)$_4$3' ssDNA. Selected classes showing streptavidin associated with a curved U- or V- shape with cartoon interpretations are indicated with red dots and expanded with cartoon interpretations (right side). **(B)** 2D-class averages of POT1/TPP1/GFP-TIN2 bound to streptavidin-5'Biotin(GGTTAG)$_4$3' ssDNA. Selected classes showing streptavidin associated with a curved U- or V- shape are indicated with red dots and expanded (right side). **(C)** 2D-class averages of the TRF1 complex bound to streptavidin-5'Biotin(GGTTAG)$_4$3' ssDNA. Selected classes showing streptavidin associated with a curved U- or V- shape are indicated with red dots and expanded (right side).

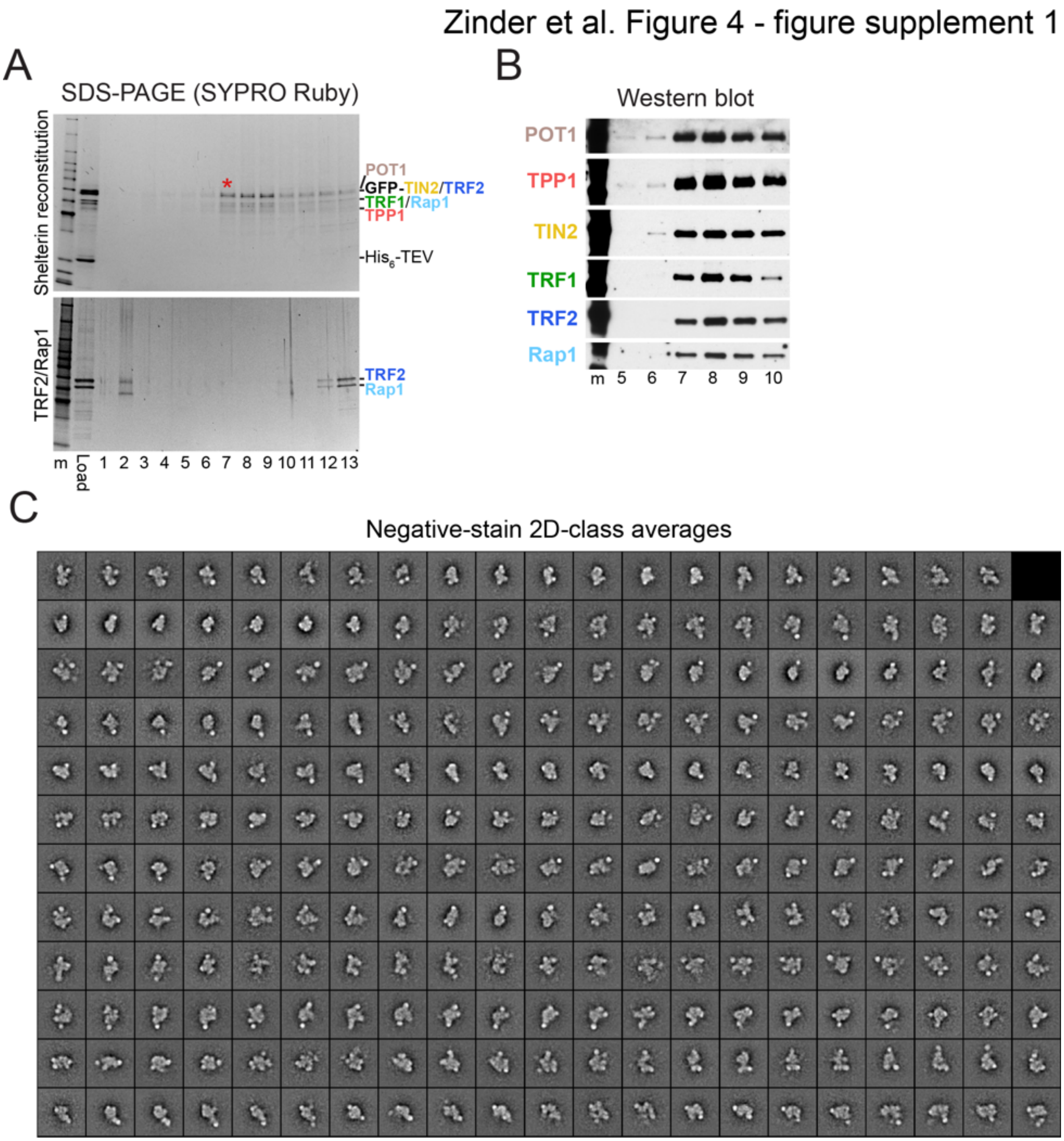


**Figure 4—figure supplement 1. Reconstitution and negative-stain EM of shelterin**
**(A)** SDS-PAGE of glycerol-gradient fractions of TRF2/Rap1 in the presence or absence of the TRF1 complex. Lower number fractions have higher glycerol concentrations. Gel is 4-12% Bis-Tris run in MOPS-SDS buffer and stained with SYPRO Ruby. Molecular-weight markers are shown in kDa. Asterisk corresponds to the fraction used for negative-stain EM analysis. **(B)** Western blot analysis of peak fractions from the glycerol gradient in panel A. Asterisk corresponds to the fraction used for negative-stain EM analysis. **(C)** Negative-stain EM 2D-class averages of reconstituted shelterin.

Zinder et al. Figure 4 - figure supplement 2

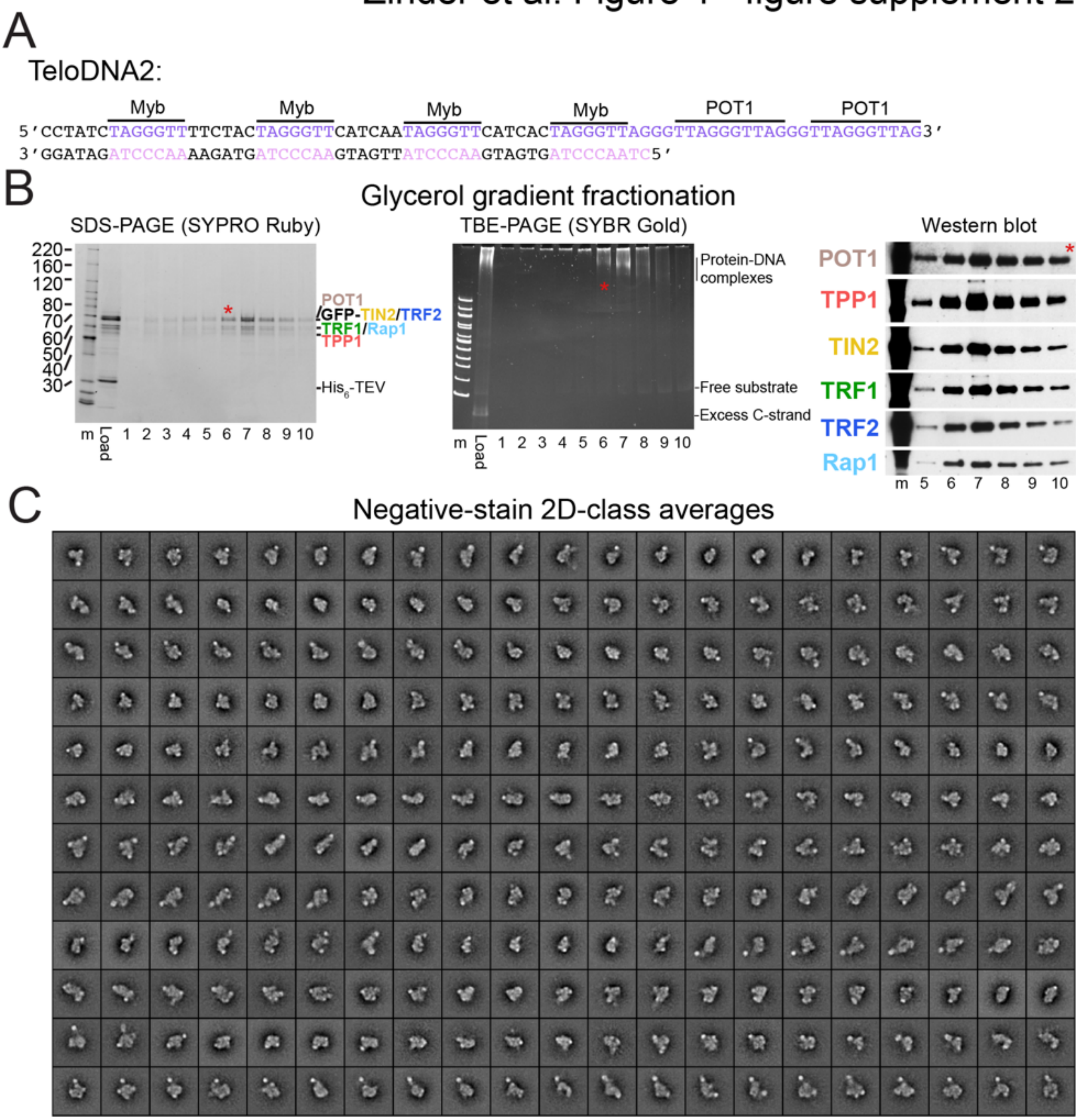


**Figure 4—figure supplement 2. Reconstitution and negative-stain EM of shelterin bound to DNA**

**(A)** DNA sequence of TeloDNA2. **(B)** SDS-PAGE, native PAGE, and Western blot of the reconstituted shelterin-TeloDNA2 complex. Lower number fractions have higher glycerol concentrations. Gel is 4-12% Bis-Tris run in MOPS-SDS buffer and stained with SYPRO Ruby for SDS-PAGE and 4-20% polyacrylamide TBE and stained with SYBR Gold for TBE-PAGE. Molecular-weight markers are shown in kDa. Asterisk corresponds to the fraction used for negative-stain EM analysis. **(C)** Negative-stain 2D-class averages of reconstituted shelterin bound to DNA.

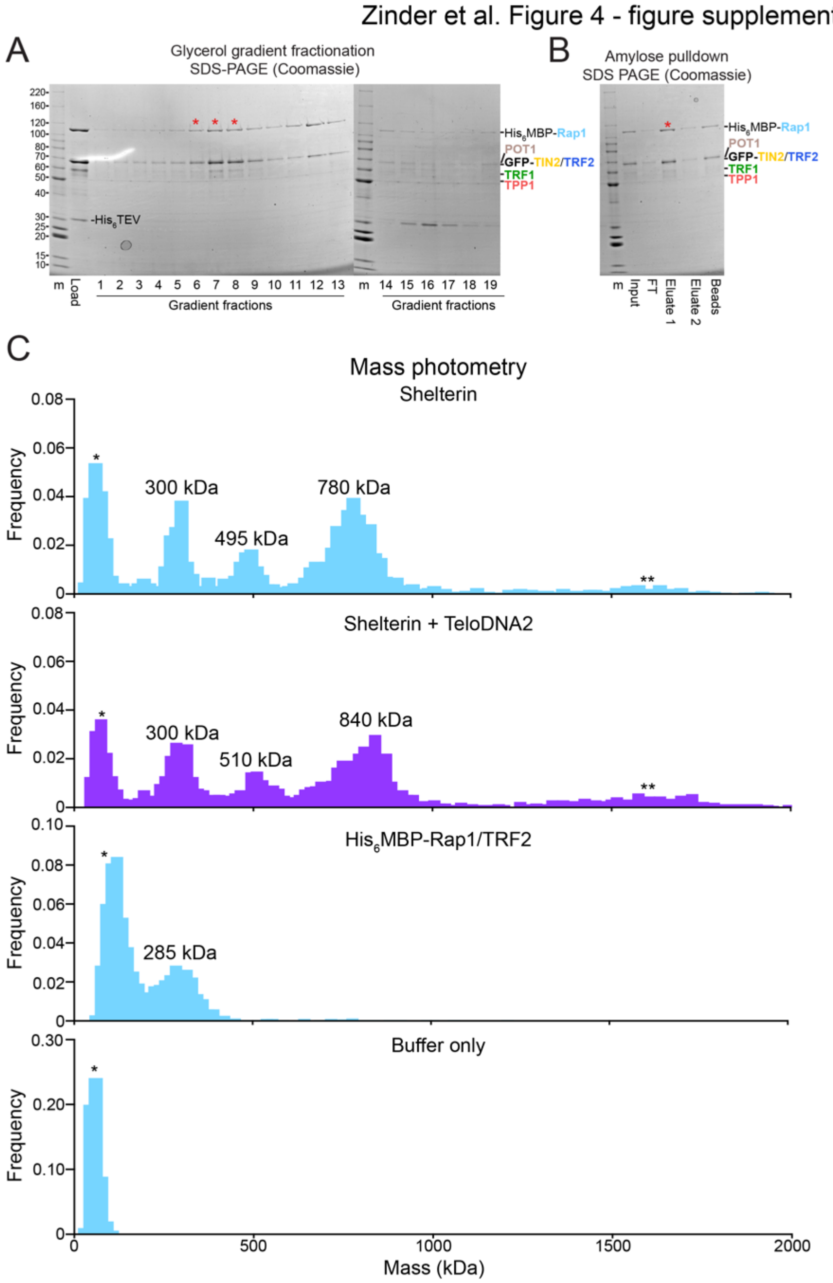


**Figure 4—figure supplement 3. Amylose pulldown and mass photometry of His6MBP-Rap1 containing shelterin**

**(A)** SDS-PAGE of glycerol-gradient fractions from the shelterin reconstitution using $His_6$MBP-Rap1. Lower number fractions have higher glycerol concentrations. Gel is 8-16% Tris-Glycine run in Tris-Glycine-SDS buffer and stained with Coomassie Blue. Molecular-weight markers (m) are shown in kDa. Red asterisks correspond to the fractions used for the amylose pulldown. **(B)** Amylose pulldown of reconstituted $His_6$MBP-Rap1-containing shelterin. Gel is 8-16% Tris-Glycine run in Tris-Glycine-SDS buffer and stained with Coomassie Blue. Red asterisk corresponds to the fraction used for mass photometry. FT, flowthrough. **(C)** Mass photometry data from Figure 4E with data at higher molecular weights included. Additionally, mass photometry measurements are shown for TRF2/$His_6$MBPRap1 and buffer alone. A single asterisk indicates a peak arising from buffer contaminants and double asterisk indicates species far outside of the instrument's calibrated range.

### Tables

**Supplementary Table 1 – Negative-stain EM Summary**

| Proteins | DNA | Figure(s) | Micrographs | Particles | Particles Per Class |
|---|---|---|---|---|---|
| **POT1/TPP1$^N$** | 5'Biotin$(GGTTAG)_2$3' | 1D, F1S1A | 100 | 40,955 | 80-200 |
| **POT1/TPP1$^N$ + streptavidin** | 5'Biotin$(GGTTAG)_2$3' | 1E, F1S1B | 75 | 24,820 | 80-200 |
| **POT1/TPP1/GFP-TIN2** | 5'Biotin$(GGTTAG)_2$3' | F2S1 | 100 | 32,813 | 60-150 |
| **POT1/TPP1/GFP-TIN2 + streptavidin** | 5'Biotin$(GGTTAG)_2$3' | 2C, F2S2A | 140 | 33,117 | 60-150 |
| **POT1$^\Delta$/TPP1$^\Delta$/GFP-TIN2$^\Delta$ + streptavidin** | 5'Biotin$(GGTTAG)_2$3' | 2C, F2S2B | 110 | 30,154 | 60-150 |
| **POT1/TPP1/GFP-TIN2/TRF1** | TeloDNA1 | 3F, F3S2A | 100 | 22,851 | 40-100 |
| **POT1/TPP1/GFP-TIN2/TRF1** | None | F3S2B | 100 | 15,058 | 40-100 |
| **POT1/TPP1$^N$ + streptavidin** | 5'Biotin$(GGTTAG)_4$3' | F3S3A | 110 | 31,399 | 40-100 |
| **POT1/TPP1/GFP-TIN2 + streptavidin** | 5'Biotin$(GGTTAG)_4$3' | F3S3B | 150 | 29,112 | 40-100 |
| **POT1/TPP1/GFP-TIN2 + TRF1 + streptavidin** | 5'Biotin$(GGTTAG)_4$3' | F3S3C | 200 | 36,202 | 40-100 |
| **Shelterin** | None | 4D, F4S1C | 300 | 31,939 | 60-150 |
| **Shelterin** | TeloDNA2 | 4D, F4S2C | 300 | 31,285 | 60-150 |

**Supplementary Table 2 – Intra-protein crosslinks within POT1/TPP1/GFP-TIN2**

| Protein | Residue 1 | Residue 2 | Lowest score | # of precursors | Interacting domains |
|---|---|---|---|---|---|
| POT1 | 427 | 433 | 2.0E-18 | 126 | HJRL |
| TIN2 | 81 | 119 | 9.5E-18 | 36 | TRFH |
| POT1 | 355 | 433 | 4.9E-30 | 12 | OB3-HJRL |
| TIN2 | 101 | 106 | 7.0E-17 | 55 | TRFH |
| POT1 | 422 | 433 | 6.0E-25 | 62 | HJRL |
| POT1 | 121 | 433 | 1.6E-21 | 6 | OB1-HJRL |
| TIN2 | 121 | 353 | 9.5E-18 | 5 | TRFH-CTD |
| POT1 | 171 | 234 | 2.7E-16 | 145 | OB2 |
| POT1 | 412 | 433 | 1.1E-15 | 14 | HJRL |

| | | | | | |
|---|---|---|---|---|---|
| POT1 | 121 | 355 | 4.0E-15 | 15 | OB1-OB3 |
| POT1 | 121 | 469 | 4.3E-15 | 10 | OB1-HJRL |
| POT1 | 433 | 469 | 5.1E-15 | 15 | HJRL |
| POT1 | 121 | 234 | 1.6E-14 | 9 | OB1-OB2 |
| POT1 | 353 | 469 | 7.6E-14 | 3 | OB3-HJRL |
| POT1 | 85 | 355 | 4.1E-13 | 5 | OB1-OB3 |
| POT1 | 234 | 469 | 7.0E-11 | 6 | OB2-HJRL |
| POT1 | 85 | 433 | 8.3E-11 | 1 | OB1-HJRL |
| POT1 | 234 | 355 | 5.1E-10 | 4 | OB2-OB3 |
| POT1 | 33 | 355 | 7.7E-10 | 7 | OB1-OB3 |
| POT1 | 234 | 433 | 1.1E-09 | 3 | OB2-HJRL |
| POT1 | 33 | 353 | 1.3E-09 | 8 | OB1-OB3 |
| POT1 | 234 | 289 | 1.5E-09 | 48 | OB2 |
| POT1 | 85 | 469 | 2.8E-09 | 8 | OB1-HJRL |
| POT1 | 121 | 121 | 3.0E-08 | 9 | OB2 |
| POT1 | 353 | 433 | 3.0E-08 | 1 | OB3-HJRL |
| POT1 | 234 | 353 | 1.4E-07 | 3 | OB2-OB3 |
| POT1 | 182 | 469 | 3.0E-07 | 5 | OB2-HJRL |
| POT1 | 355 | 379 | 3.6E-06 | 6 | OB3 |
| POT1 | 131 | 353 | 3.7E-06 | 4 | OB1-OB3 |
| POT1 | 407 | 433 | 7.0E-06 | 6 | HJRL |
| POT1 | 379 | 433 | 3.1E-05 | 3 | OB3-HJRL |
| TIN2 | 233 | 235 | 3.4E-05 | 3 | CTD |
| TIN2 | 62 | 81 | 1.5E-04 | 10 | TRFH |
| POT1 | 469 | 504 | 2.1E-04 | 5 | HJRL |
| POT1 | 131 | 469 | 2.5E-04 | 4 | OB1-HJRL |
| POT1 | 85 | 131 | 4.1E-04 | 1 | OB2 |
| POT1 | 131 | 234 | 5.6E-04 | 7 | OB1-OB2 |
| TIN2 | 101 | 119 | 6.1E-04 | 3 | TRFH |
| POT1 | 353 | 355 | 6.6E-04 | 1 | OB3 |
| POT1 | 379 | 469 | 8.0E-04 | 4 | OB3-HJRL |
| POT1 | 469 | 469 | 8.8E-04 | 2 | HJRL |
| POT1 | 355 | 469 | 1.4E-03 | 1 | OB3-HJRL |
| POT1 | 422 | 469 | 1.9E-03 | 8 | HJRL |
| POT1 | 131 | 355 | 2.8E-03 | 6 | OB1-OB3 |

**Supplementary Table 3 – Inter-protein crosslinks within POT1/TPP1/GFP-TIN2**

| Protein1 | Residue | Protein 2 | Residue | Lowest score | # of precursors | Interacting domains |
|---|---|---|---|---|---|---|
| POT1 | 433 | TPP1 | 232 | 1.2E-20 | 19 | HJRL-OB |
| POT1 | 433 | TPP1 | 170 | 5.2E-21 | 19 | HJRL-OB |
| POT1 | 469 | TPP1 | 232 | 1.7E-07 | 11 | HJRL-OB |
| POT1 | 121 | TIN2 | 81 | 2.0E-14 | 10 | OB1-TRFH |
| POT1 | 469 | TPP1 | 170 | 2.2E-12 | 9 | HJRL-OB |
| POT1 | 234 | TIN2 | 81 | 3.3E-08 | 8 | OB2-TRFH |
| TIN2 | 81 | TPP1 | 170 | 2.1E-06 | 7 | TRFH-OB |
| POT1 | 433 | TIN2 | 101 | 9.3E-15 | 6 | HJRL-TRFH |
| POT1 | 121 | TIN2 | 98 | 1.3E-06 | 6 | OB1-TRFH |
| POT1 | 469 | TPP1 | 233 | 2.8E-11 | 5 | HJRL-OB |
| POT1 | 85 | TPP1 | 492 | 3.0E-08 | 5 | OB1-TIN2BD |
| POT1 | 355 | TPP1 | 492 | 3.9E-05 | 5 | OB3-TIN2BD |
| POT1 | 234 | TIN2 | 131 | 4.6E-04 | 5 | OB2-TRFH |
| POT1 | 234 | TPP1 | 170 | 2.3E-13 | 4 | HJRL-OB |
| POT1 | 353 | TPP1 | 170 | 6.2E-12 | 4 | HJRL-OB |
| POT1 | 355 | TIN2 | 101 | 7.9E-07 | 4 | OB3-TRFH |
| POT1 | 234 | TIN2 | 106 | 1.9E-06 | 4 | OB2-TRFH |
| POT1 | 234 | TIN2 | 101 | 3.0E-05 | 4 | OB2-TRFH |
| POT1 | 469 | TIN2 | 106 | 2.3E-15 | 3 | HJRL-TRFH |
| POT1 | 85 | TPP1 | 170 | 9.6E-14 | 3 | OB1-OB |
| POT1 | 469 | TPP1 | 492 | 2.6E-06 | 3 | HJRL-TIN2BD |
| POT1 | 430 | TIN2 | 101 | 6.4E-11 | 2 | HJRL-TRFH |
| POT1 | 433 | TPP1 | 492 | 3.7E-10 | 1 | HJRL-TIN2BD |
| POT1 | 234 | TIN2 | 119 | 2.4E-07 | 1 | OB2-TRFH |
| POT1 | 85 | TIN2 | 101 | 2.0E-06 | 1 | OB1-TRFH |
| POT1 | 433 | TIN2 | 98 | 9.9E-06 | 1 | HJRL-TRFH |
| POT1 | 234 | TIN2 | 233 | 4.4E-05 | 1 | OB2-CTD |
| POT1 | 469 | TIN2 | 235 | 4.4E-04 | 1 | HJRL-CTD |
| POT1 | 427 | TPP1 | 170 | 9.4E-04 | 1 | HJRL-OB |
| POT1 | 353 | TPP1 | 492 | 1.3E-03 | 1 | OB3-TIN2BD |
| POT1 | 433 | TIN2 | 81 | 2.8E-03 | 1 | HJRL-TRFH |

**Supplementary Table 4 – Predicted and measured molecular weights in kDa of shelterin and subcomplexes**

| Complex | Predicted Molecular weight | Mass Photometry Measured | Native Mass Spectrometry Measured |
|---|---|---|---|
| **$(TRF1)_2$** | 97.057 | 80-90 | 97.070 |
| **Twinstrep-POT1/TPP1/GFP-TIN2** | 191.578 | 170-210 | 191.623, 191.659, 191.908 |
| **Twinstrep-POT1/TPP1/GFP-TIN2/$(TRF1)_2$** | 288.635 | 270-290 | 288.806 |
| **$(Twinstrep\text{-}POT1/TPP1/GFP\text{-}TIN2/TRF1)_2$** | 480.213 | 470-490 | 480.700 |
| **$(TRF1)_2$/GFP-TIN2** | 164.276 | N/A | 164.325 |
| **Twinstrep-POT1/TPP1** | 124.359 | N/A | 124.472 |
| **Twinstrep-POT1** | 75.438 | N/A | 75.504 |
| **$(TRF2/His_6MBP\text{-}Rap1)_2$** | 286.420 | 280-300 | N/A |
| **POT1/TPP1/GFP-TIN2/$(TRF2/His_6MBP\text{-}Rap1)_2$** | 474.13 | 495? | N/A |
| **$(POT1/TPP1/GFP\text{-}TIN2/TRF1/TRF2/His_6MBP\text{-}Rap1)_2$** | 758.568 | 780 | N/A |
